# m6AConquer: a Data Resource for Unified Quantification and Integration of m6A Detection Techniques

**DOI:** 10.1101/2024.09.10.612173

**Authors:** Xichen Zhao, Haokai Ye, Tenglong Li, Daniel J. Rigden, Zhen Wei

## Abstract

N6-methyladenosine (m^6^A) is the most prevalent RNA modification in mammalian cells and the most extensively studied epitranscriptomic mark. More than 10 m^6^A detection techniques have been proposed to measure m^6^A stoichiometry at either the site or peak level. However, these detection techniques are processed through heterogeneous pipelines, using different computational filters and reference features, leading to difficulties in fully harnessing data integration and analysis across orthogonal m^6^A detection techniques. Our m6AConquer (**Con**sistent **Qu**antification of **E**xternal **m**^**6**^**A R**NA Modification Data) tackles this challenge by establishing a consistent multi-omics data-sharing standard, summarizing quantitative m^6^A data from 10 detection techniques using a unified reference feature set. Furthermore, we standardize site calling and m^6^A count matrix normalization procedures across platforms through a computational framework that accounts for over-dispersion in m^6^A levels. Available m^6^A detection techniques can be categorized into four types: antibody-assisted, chemical-assisted, enzyme-assisted, and direct-RNA sequencing. We leverage this categorization to develop a reproducibility-based integration framework that enables the reliable detection of high-confidence m^6^A sites confirmed across orthogonal techniques. Empirical evaluations report that both the site-calling and the integration framework outperform common alternatives, enhancing biological relevance. We apply interpretable machine learning models on our integrated high-confidence sites, and the result consistently identify proximity to intron-exon junctions as the driving predictor of m^6^A site coordinates across different techniques, demonstrating the high quality of the data curated in m6AConquer. m6AConquer webserver is freely accessible under: http://rnamd.org/m6aconquer.

**GRAPHICAL ABSTRACT:** 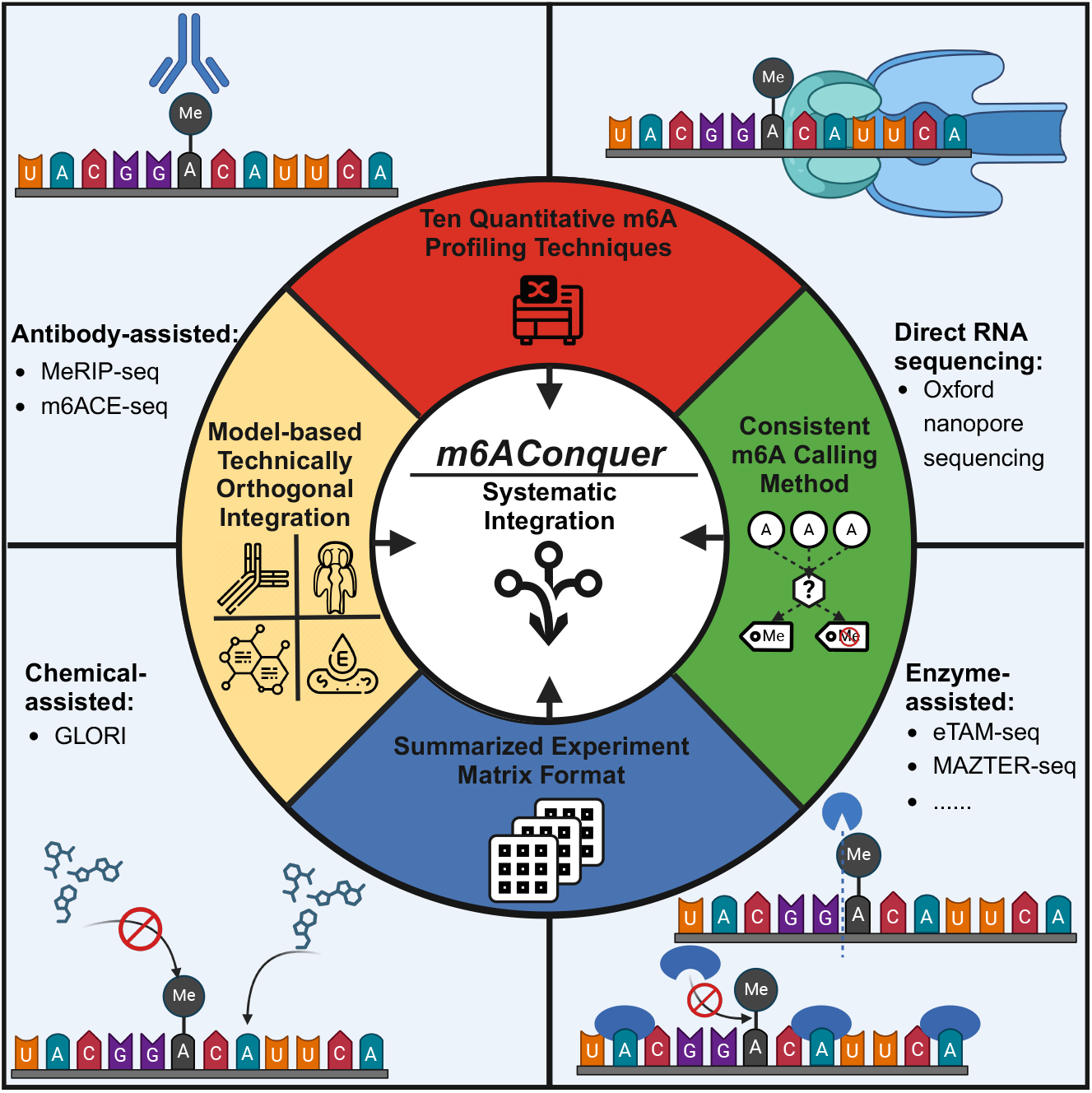

## INTRODUCTION

N6-methyladenosine (m^6^A) modification is one of the most widely studied RNA chemical modifications in eukaryotic cells. m^6^A epitranscriptomically regulates gene expression via specifying RNA half-life (1-3). It is the most abundant type of RNA modification, topologically and functionally similar to DNA m5C modification. The basal constitutional level of m^6^A on transcripts functions as a negative gene expression regulation mechanism by targeting the modified RNA to P-bodies for degradation (4-8). The installation of m^6^A on transcripts is mainly dictated by the DARCH consensus motif, but it is suppressed in regions close to splicing junctions (9). This mechanism makes the intron-exon architecture a key determinant of RNA half-life, with longer exons harbouring more m^6^A modifications (3,10,11).

m^6^A modification state is dynamically regulated by protein factors, including writers (METTL3, METTL14, WTAP, KIAA1429), erasers (FTO, ALKBH6), and readers (YTHDF1-3, YTHDC1-2, HNRNPC) (12,13). The specific m^6^A regulators allow the same m^6^A site link to distinct downstream outcomes via alternative mechanisms (2,4,14-20). m^6^A can also interact with epigenetic modules, including histone methylation (H3K36me3) and transposable element (SINEs and LINEs) expression (21-27). m^6^A on noncoding RNAs, such as miRNAs and carRNAs, has also been reported to influence their processing and function. Furthermore, m^6^A plays a crucial role in fundamental cellular processes such as the cell cycle, differentiation, and stress responses (4,28-33). Accordingly, m^6^A is known to be involved in development of disease and other medical conditions, including acute myelogenous leukemia (AML) (34-36), depression (37-39), and multiple cancers (12,15,40,41). Perturbation of m^6^A regulators, like FTO, can also impact crop yields, as observed in maize (42-44).

Existing sequencing techniques to measure m^6^A methylome can be broadly classified into four categories: antibody-assisted, enzyme-assisted, direct-RNA sequencing, or chemical-assisted. In the antibody-assisted approach, anti-m^6^A antibodies recognize m^6^A sites followed by sequencing (45-47). Two best-known examples of the antibody-assisted approach are MeRIP-seq (45) and miCLIP. The enzyme-assisted approach involves inducing covalent changes at m^6^A bases using enzymes or other protein factors. Examples are enzyme conversion-based eTAM-seq (48) and restriction enzyme-based MAZTER-seq (49). The ONT direct-RNA sequencing approach can recognize unique alterations in ionic current as modified nucleic acids pass through a nanopore protein (50-53). Finally, the chemical-assisted approach of GLORI (54) relies on a chemical transformation reaction of unmethylated Adenine to Inosine, which is conceptually similar to bisulfite sequencing for DNA m^5^C.

Some techniques for m^6^A profiling are absolute quantification methods; others are not. Absolute quantification technologies, such as GLORI and eTAM-seq, have the advantage of providing precise binomial data at read level. This includes the exact counts of m^6^A-modified reads and total read coverage at specific sites. In contrast, antibody-dependent techniques like MeRIP-seq and m6ACE-seq (46,47) rely on the ratio of two-sample data from immunoprecipitated (IP) and input. However, the absolute sequencing depth difference between the samples is unidentifiable. Consequently, leveraging absolute quantification data that eliminate technical confounding factors is crucial for the advancement of epitranscriptomics.

Nevertheless, each of the four categories of m^6^A detection strategies possesses intrinsic limitations. Around 30% of peaks identified by the antibody-assisted technique of MeRIP-seq are false positives due to non-specific antibody binding (59,60,91). These false positives can be due to regions with similar motif sequences to true m^6^A sites. The off-target sites are typically located in high-coverage areas of short exons and transcripts, complicating biological data analysis (55,56). Artifacts in enzyme-assisted techniques depend on the protein used. For example, the restriction enzyme-based MAZTER-seq can have up to 50% false positives due to inaccurate cleavage (54,56). The precision of direct RNA sequencing is also limited because of the overlap between modified and background signals. Additionally, the accuracy is further affected by the incomplete sequence contexts in its IVT training data (56,57). Finally, chemical-assisted techniques can suffer from incomplete conversion, as seen in bisulfite sequencing, often due to the secondary structure of RNA(58,59).

To this end, it is crucial to remove technique-specific artifacts using integrative data from technically independent measurements. In recent studies, technically orthogonal validations have been increasingly adopted, where intersections between techniques reinforce the understanding of true m^6^A installation mechanism in the transcriptome (3,55,60,61). For example, using common sites from miCLIP, m6A-SAC-seq, DART-seq, and GLORI, Anna *et al*. show that m^6^A formation is driven by the splice-site exclusion mechanism (3,10). Another study using a similar integrative analysis found that m^6^A does not promote protein translation (61).

To address the difficulties of cross-modality m^6^A detection, we here introduce a new database m6AConquer which implements a model-based, technical independent integration approach to enhance the fidelity of integration across heterogeneous datasets and platforms. Although published works use multiple m^6^A profiling techniques to validate their conclusions, they do not apply a statistical framework to measure reproducibility and eliminate spurious overlaps. This work attempts to fill this gap by introducing irreproducible discovery rate (IDR) integration for orthogonal techniques in m^6^A data. IDR is well-established in DNA epigenetics and used by the ENCODE database (62) to integrate CHIP-Seq samples (63,64). The IDR model effectively uses rank information to quantify uncertainties in overlaps between replicating omics experiments. The reproducible sites called by IDR are both highly ranked and positively correlated between paired omics assays.

Furthermore, m6AConquer collects absolute quantification m^6^A data and transforms samples from all techniques into a standard matrix format, in which all datasets use consistent m^6^A sites as reference features. This effort aims to address a gap in established m6Aomics resources like m6A-Atlas2 (65), MeT-DB2 (66), RMBase2 (67), and DirectRMDB (68), which lack a consistent and efficient data framework for m^6^A data sharing (Table 1). All these databases gather sites or peaks returned by different site-calling pipelines, each with different statistical assumptions and filtering criteria. This results in inconsistent site features due to varying thresholds applied after site calling. In addition, to the best of our knowledge, none of these databases have collected absolute quantification data from GLORI and eTAM-seq. For example, m6A-Atlas2 only quantifies MeRIP-seq using IP over input fold change, while other techniques gathered are presented as sites or peaks. Moreover, the m6AConquer data-sharing framework further extends to the multi-omics level (combining m^6^A-omics with transcriptomics). This new feature enables efficient multi-omics data analysis, as has been achieved in the scRNA-seq field with GTEx (69) and conquer (70).

**Table 1.**
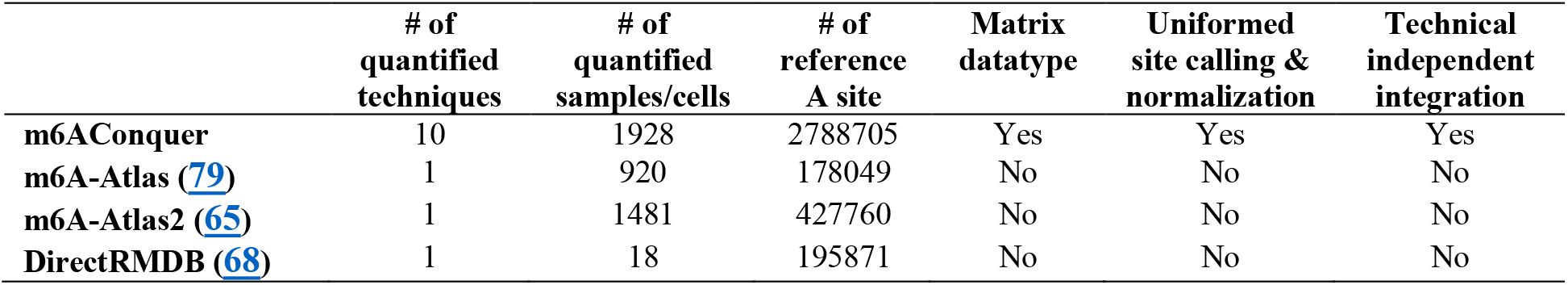
Comparison of features across m6AConquer, m6A-Atlas, m6A-Atlas2, and Direct-RMDB databases for the human genome.

On top of that, m6AConquer identifies a robust site-calling and normalization method, applying it uniformly across different m^6^A detection platforms. Various statistical testing methods have been developed to call m^6^A sites in different detection techniques. For example, GLORI and eTAM-seq use a single binomial distribution to calculate p-values (48,54). While Poisson distribution-based tests are employed in MeRIP-seq peak calling with MACS2 (71,72) and exomePeak (73). More advanced statistical models, such as the beta-binomial distribution, are implemented in the MeTPeak (74). The beta-binomial distribution accounts for over-dispersion in m^6^A ratios making it a strong candidate for a general cross-platform site caller. Beta-binomial distribution-based models have also been used for site calling and cross-platform normalization in DNA m5C studies (75-77).

Beyond that, the integrative omics database of m6AConquer can also support faithful and large-scale machine learning (ML) modeling. Batch effects and platform-specific artifacts can introduce significant obstacles to ML imputation on heterogeneous datasets. The ENCODE imputation challenge project (78) highlighted that ML evaluations are often confounded by distributional shifts due to differences in data collection and processing steps. Therefore, the novel features introduced by m6AConquer—such as the consistent matrix-based omics data framework, uniform site-calling and normalization procedures, and model-based integration that accounts for technical independence—are essential for obtaining reliable biological insights on m^6^A using ML.

## MATERIAL AND METHODS

### Processing and Calibration of m^6^A Profiling Techniques

Raw sequencing data from 10 quantitative m^6^A profiling techniques, including antibody-assisted (MeRIP-seq (65), m6ACE-seq (46,47)), enzyme-assisted (eTAM-seq (48), m6A-SAC-seq (80,81), MAZTER-seq (49), DART-seq (82), scDART-seq (83), and m6A-REF-seq (56,84)), Direct-RNA Sequencing (53), and chemical-assisted (GLORI (54)) methods, were collected and processed. A total of 1,928 FASTQ and FAST5 datasets were retrieved from GEO (85,86), GSA (87,88), and EMBL-EBI (89), then processed and quality controlled using the respective pipelines in the references. Hisat2 and STAR were used for regular RNA-Seq alignment, Hisat3N for nucleotide conversion-based techniques, and Minimap2 for DRS reads. Reference genomes hg38/GRCh38 (human) and mm10/GRCm38 (mouse) were used. Read quality control was performed with Cutadapt, Fastp, and UMItools, as specified by the original studies.

In m6AConquer, data from diverse platforms were consistently quantified against a unified reference feature set to ensure integration. For the human m^6^A methylome, 2,526,220 DRACH motifs on exons from the UCSC hg38 annotation served as the primary reference. Additionally, 262,485 GLORI-identified m^6^A sites from Cong’s study (54) that did not map to exonic DRACH motifs were included to capture non-DRACH and intronic m^6^A sites. This consistent feature set was used to quantify all 1,928 human samples or cells at base resolution across ten techniques.

To maintain raw read count integrity at reference sites, no premature filtering by site calling was performed post-alignment. Alignment results were counted against consistent reference sites to generate m6A counts and total read coverage using Python scripts. For non-absolute quantification techniques of MeRIP-seq and m6ACE-seq, m^6^A counts were derived from counting overlapped IP read fragments, while total coverage was the sum of the count in IP and the paired input control samples. Figure 1 presents the m6AConquer workflow, with data collection and pipeline configurations detailed in the Supplementary Material.

**Figure 1.**
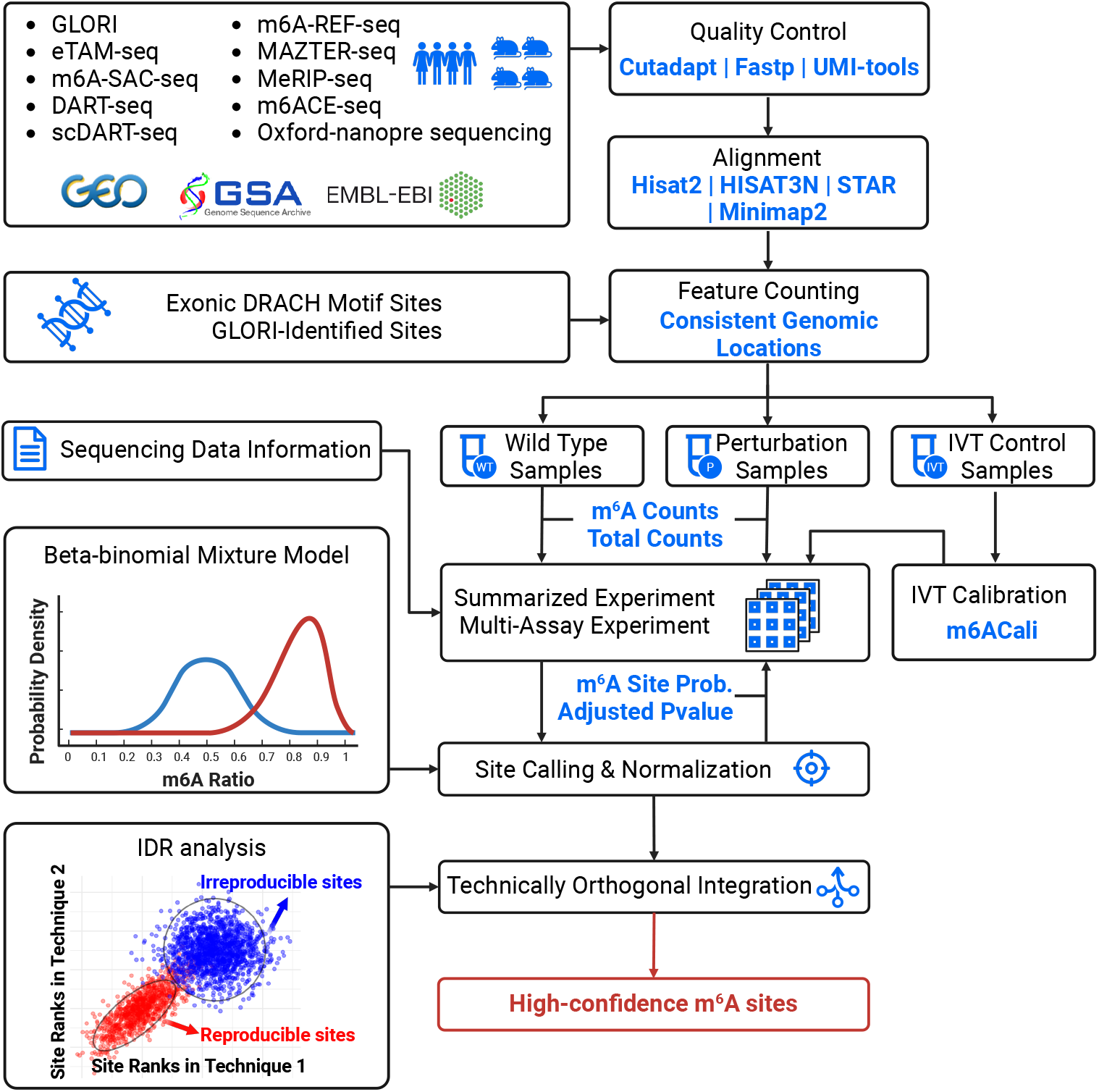
The m6AConquer pipeline integrates m^6^A data from 10 detection techniques in human and mouse. Raw reads are first quality controlled (Cutadapt, Fastp, UMI-tools) and aligned (STAR, Hisat2, Hisat3N, Minimap2) to genome reference. Feature counting is then consistently applied to exonic DRACH motifs and GLORI-identified sites across all datasets. Samples, categorized as wild type, perturbation, or IVT controls, are processed for site calling (adjusted p-values) and normalization (m^6^A site probabilities) using a beta-binomial mixture model. IVT controls are used to calibrate regular samples, with MeRIP-Seq calibrated using m6ACali. Technically orthogonal integration is performed using IDR analysis on wild-type samples to identify high-confidence m^6^A sites.

For MeRIP-seq, GLORI, and eTAM-seq, high-quality in-vitro transcribed (IVT) experiments (60, 54 48) were used as negative controls to calibrate read count matrices by masking false positive sites. The false positive sites, identified in IVT samples, were set to zero in the count matrices for both m^6^A and total read coverage. For MeRIP-seq, false positives were detected using the machine learning-powered m6ACali method (55) with a probability threshold of 0.5. For GLORI and eTAM-seq, IVT samples were quantified against reference sites as the regular samples, and false positives were defined by an adjusted p-value below 0.05 after BBmix site calling.

### Standardized Data Sharing Framework in m6AConquer

To maximize the use of matrix-based quantification data, single-omics and multi-omics data frameworks were assembled using Bioconductor objects. For the single-omics data framework, SummarizedExperiment (SE) objects were created by combining multiple genomic assays (matrices) with matching dimensions (see Figure 2A). The m^6^A-omics SE includes four assays: m^6^A count, total coverage, m^6^A foreground probability, and BH-adjusted p-value. RowRanges are set to the GenomicRanges of reference m^6^A sites, while the column table contains sample-specific annotations, including accessions, conditions, perturbations, and techniques. Metadata encompasses site-calling parameters for each sample and high-confidence integrated sites identified via IDR.

**Figure 2.**
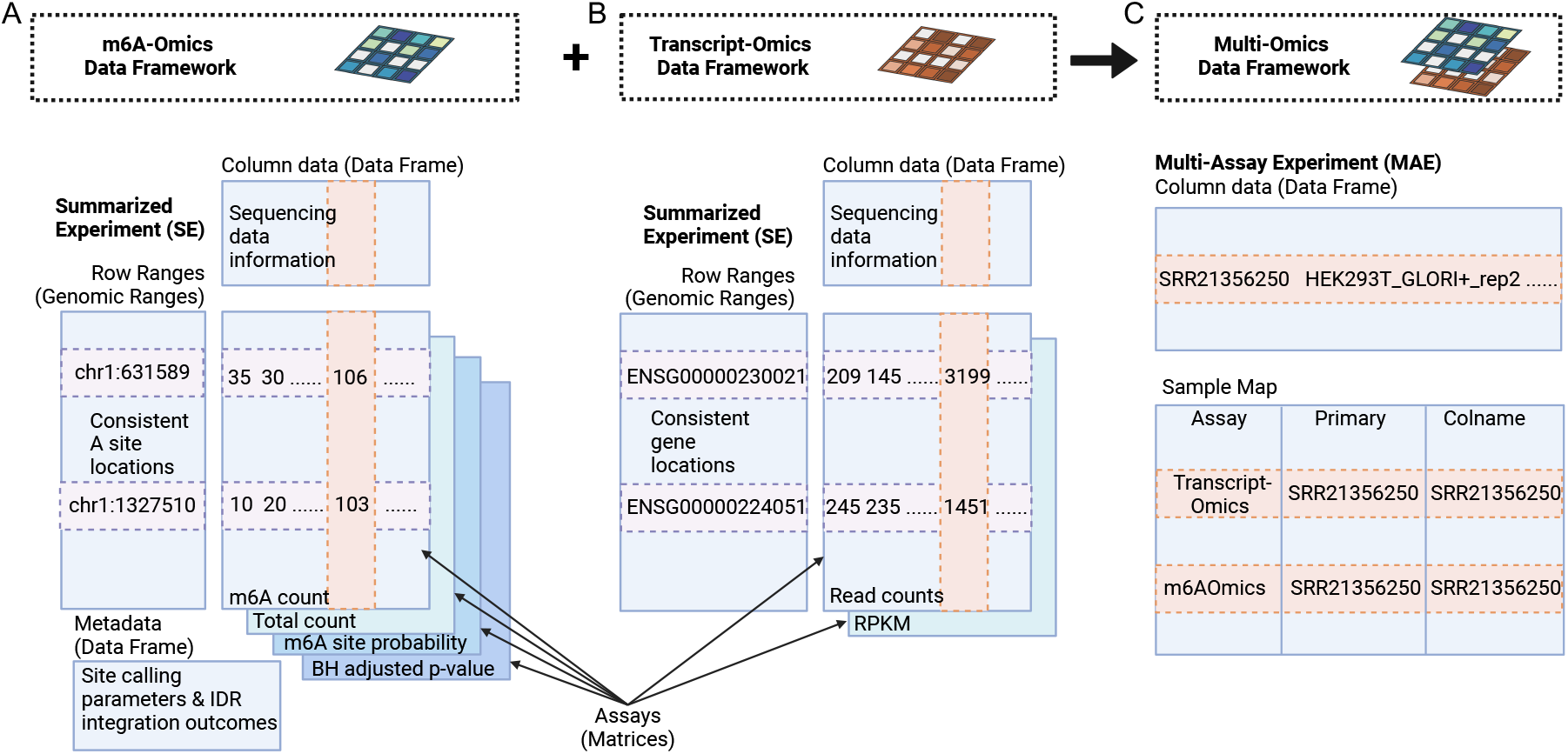
The consistent data sharing frameworks established in m6AConquer. (**A**) m6A-Omics Data Framework organizes m^6^A quantification data into a SummarizedExperiment (SE) object, including reference A site locations (RowRanges), sequencing information (Column Data), and matrices (Assays) for m^6^A counts, total coverage, m6A site probabilities, and p-values. Metadata covers site calling parameters and IDR outcomes. (**B**) Transcript-Omics Data Framework contains a similar SE object for transcriptomic (expression) data, including gene features, sequencing information, and matrices for read counts and RPKM values. (**C**) Multi-Omics Data Framework integrates m^6^A-omics and transcriptomics into a MultiAssayExperiment (MAE) object, combining omics modalities and a Sample Map linking assays to samples, enabling comprehensive multi-omics analysis.

Subsequently, to enable multi-omics analysis of m^6^A quantification and gene expression, we built a multi-omics data framework using the MultiAssayExperiment package (90). RNA expression levels were quantified for each m^6^A sample, with only input samples counted in antibody-assisted techniques to avoid confounding by m^6^A levels. The transcriptome data were also organized into SE objects with 58,650 gene features from GRCh38 and matrices representing read counts and RPKM values (Figure 2B). Read counts were obtained using the SummarizedOverlaps function in GenomicAlignments. The transcriptome SE columns and annotations were matched one-to-one with those in the m^6^A-omics SE. The MultiAssayExperiment (MAE) object was then created by integrating m^6^A-omics and transcriptome data (Figure 2C). All data frameworks (m^6^A-omics, transcriptomics, and multi-omics) are available for download at http://rnamd.org/m6aconquer/download.html.

### Uniform Site Calling and Normalization across m^6^A Detection Platforms

After assembling read counts in the standardized matrix format, a two-component beta-binomial mixture (BBmix) model is applied uniformly for site calling and normalization across different m^6^A quantification techniques. BBmix was originally introduced by MeTPeak (74) for peak calling in MeRIP-Seq and later used in the m^6^A database MeT-DB2 (66). The BBmix model in m^6^AConquer has been adapted from MeTPeak by dropping the HMM component to better fit reference site-based analysis. BBmix is also widely used in DNA m5C analysis (75,76), cross-platform DNA methylation normalization (77), and allele-specific expression analysis in RNA-Seq data (91,92)

For a given sample (column) of quantitative m^6^A data in the SummarizedExperiment object, let *x*_*i*_ be the read count of m^6^A at site *i*, then the observed probability distribution of *x*_*i*_ is expressed by the BBmix model using equation (1).

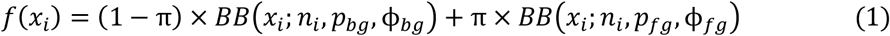

Here, *n*_*i*_ indicates the total count of read coverage at site *i, π* represents the proportion of the foreground component, and *BB* denotes the beta-binomial distribution function. *p* and *ϕ* are mean and over-dispersion parameters for the beta-binomial distributions, with “*fg*” and “*bg*” denoting the methylated m^6^A site (foreground) and unmethylated A site (background), respectively. The beta-binomial distribution enables over-dispersion in binomial distributed data, parallel to the evolution from Poisson to negative binomial distribution (93,94). Please refer to (81) for details about the parameterization on over-dispersion.

Parameters in BBmix were estimated in R using the Expectation Maximization (EM) algorithm. The beta-binomial distribution’s maximum likelihood estimation (MLE) was implemented via the Newton-Raphson method, with initial parameters set using the Method of Moments in the first EM iteration, as in prior studies (74,77,95). We also considered the zero-inflated beta-binomial mixture model (ZIBBmix), proposed for scDNA m5C analysis (95). However, its zero component weights were consistently estimated as zero across most platforms, rendering it redundant with BBmix and thus not applied in m6AConquer.

The p-value is calculated as described by MeTPeak (74), using the estimated parameters of the background beta-binomial distribution. The one-sided p-value for site calling represents the probability of observing a higher m^6^A ratio than expected based on the background distribution, as defined in equation (2).

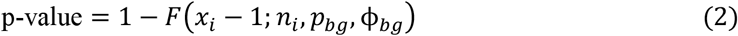

Where *F* denotes the CDF of the beta-binomial distribution, which is implemented with the *pbbinom* function in the *extraDistr* R package. Subsequently, the adjusted p-value is computed using the Benjamini-Hochberg (BH) approach implemented in the *p*.*adjust* R function.

The m^6^A site probability, m6A prob, defined in equation (3), is calculated as the posterior probability that a site belongs to the m^6^A (foreground) component. m6A prob normalizes m^6^A levels across samples and techniques by accounting for platform-specific variations, and is subsequently used as input for cross-technique integration via IDR.

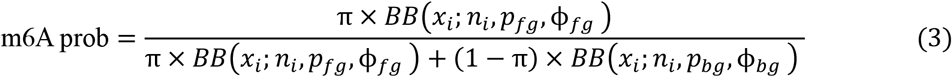

Subsequently, to justify the choice of the site-calling statistical method, the empirical model fit across various statistical models was benchmarked. Many existing pipelines rely on a single binomial distribution for site-calling from quantitative m^6^A data (48,54). Therefore, several alternative models were compared, including the single binomial, two-component binomial mixture, single beta-binomial, and beta-uniform mixture models. Bayesian Information Criterion (BIC), as defined in equation (4), was used to evaluate the model quality.

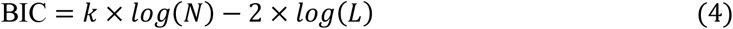

Here, *k* denotes the number of model parameters, *log* is the natural logarithm, *N* denotes the number of observations (m^6^A sites), and *L* denotes the likelihood value estimated at MLE. BIC balances goodness of fit with model complexity to prevent overfitting, making it a robust criterion for selecting biologically meaningful methods. A lower BIC indicates a more suitable model for the dataset.

BBmix and the benchmarked alternative methods were also implemented in a reproducible R package *OmixM6A*, accessible at https://github.com/ZW-xjtlu/OmixM6A. Users can download the summarized count matrix from m6AConquer and easily fit alternative p-value and m^6^A site probability using the user-friendly R package. Detailed BBmix site calling algorithm information is available in the Supplementary Material.

### Reproducibility-based m^6^A Data Integration among Orthogonal Techniques

In m6AConquer, the irreproducible discovery rate (IDR) method was used to integrate quantitative m^6^A. The IDR method for omics data integration was first proposed by Qunhua et al. (64). It is now the standard pipeline for integrating replicates in the ENCODE project (62). The IDR model converts paired omics signals from two samples into ranks, which are then integrated using a two-component bivariate Gaussian copula mixture model. Reproducible m^6^A sites are identified by positive correlation and consistently high ranks between replicates, while irreproducible m^6^A sites lack correlation and show inconsistent ranks

Specifically, the IDR value is defined by the posterior probability (responsibility) of the irreproducible component, as specified by equations (5-7).

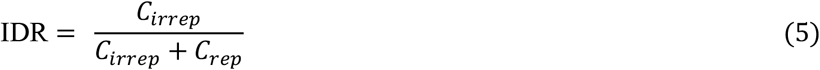

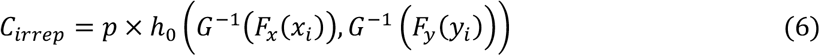

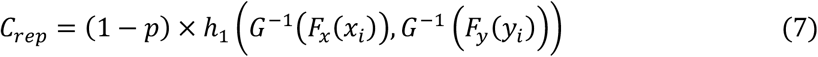

Here, 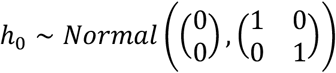 and 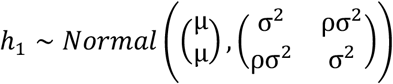. *σ*^*2*^ and *ρ* are variance and correlation parameters for the reproducible component, respectively, with *ρ* > 0. *p* specifies the proportion for the irreproducible component. Variables *x* and *y* are continuous signals (m6A prob by default) measured by two types of m6A profiling techniques respectively at site *i. F*_*x*_ and *F*_*y*_ are their empirical marginal CDFs used to compute ranks. *G*^−1^is a transformation applied in Gaussian copula modeling (see Supplementary Material or section 2.22 in (64) for details). IDR was implemented in m6AConquer using the default settings of the *idr1d* function in the Bioconductor package *idr2d*, leveraging its optimized parameter initialization for epigenetics analysis.

In the m6AConquer database, IDR is calculated between all pairs of technically independent m6A detection techniques. The technical independence is defined when two methods come from different groups in the four categories: antibody-assisted, enzyme-assisted, chemical-assisted, and direct-RNA sequencing (see Figures 3A and 3C). An exception is made for direct-RNA sequencing and antibody-assisted methods, which are also treated as technically dependent. This is because the m6Anet tool (53) used to process DRS data was trained on m6ACE-seq. High-confidence integrated sites in m6AConquer were filtered with IDR < 0.05 and adjusted p-value < 0.05 in both techniques. The high-confidence sites can be downloaded from the m6AConquer server’s download page and are also annotated in the row metadata within the SummarizedExperiment data-sharing object.

**Figure 3.**
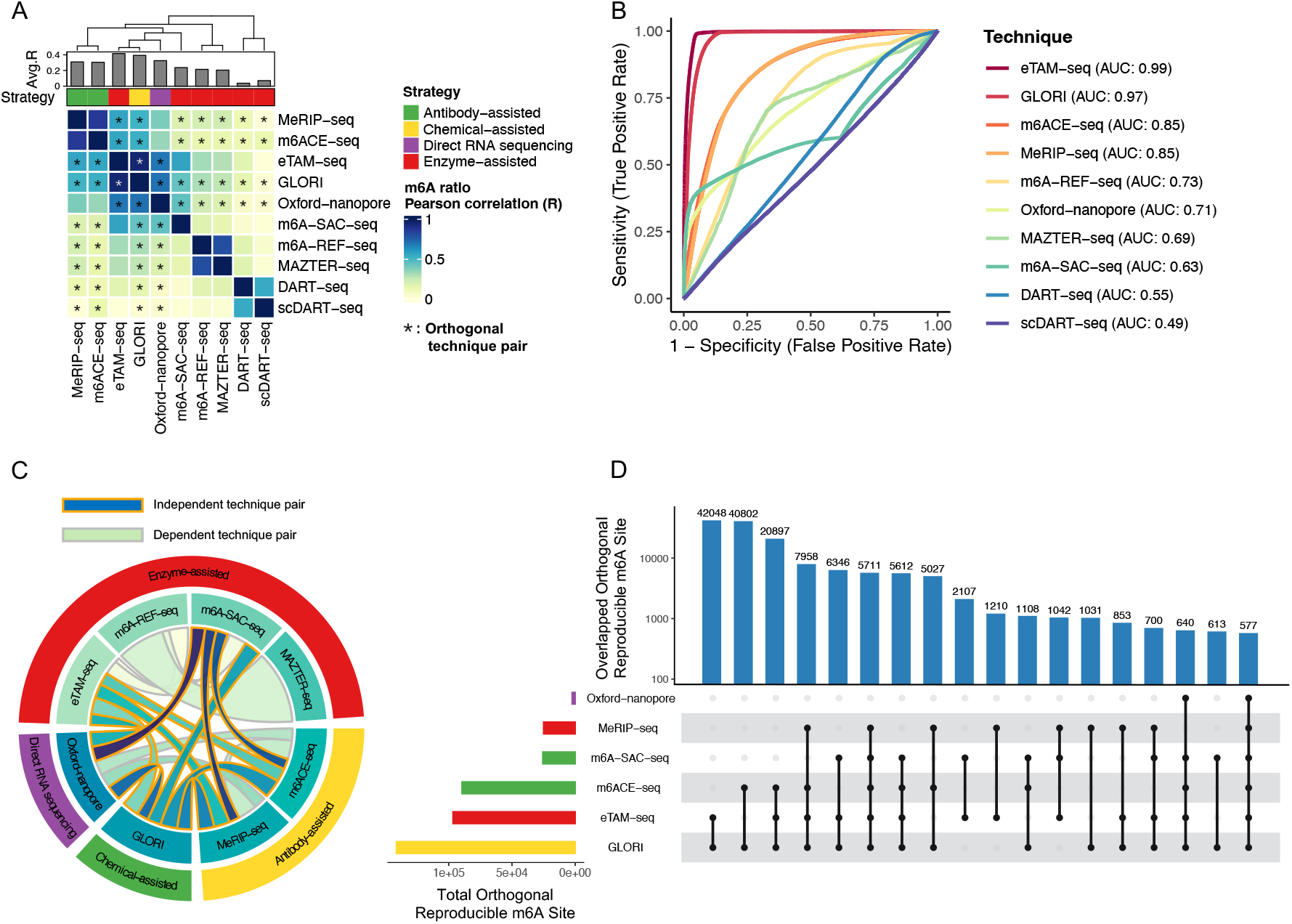
Comparative Analysis of m^6^A Detection Techniques and Integration Results (**A**) Heatmap of Pearson correlation coefficients for m^6^A methylation ratios across ten detection techniques, categorized into four groups. Orthogonal technique pairs are marked with an asterisk. (**B**) ROC curves for m^6^A site prediction accuracy using a BBmix-based Bayes classifier. GLORI (AUC = 0.97) and eTAM-seq (AUC = 0.99) show superior performance. The ground truth used for evaluation is the orthogonal reproducible sites (high-confidence integrated sites). (**C**) Circular plot showing the distribution of reproducible m^6^A sites (IDR < 0.05) identified by orthogonal (blue bands) and dependent technique pairs (green bands). The blue bands indicate the high-confidence integrated sites in m6AConquer. (**D**) Upset plot displaying the overlap of orthogonal reproducible m^6^A sites across top technique combinations, with the largest overlaps observed between GLORI and eTAM-seq, validating their reliability.

To integrate multiple samples within each technique, individual samples were pooled by summing m^6^A and total coverage counts across replicates, creating combined samples. This approach was chosen because most variance in m^6^A methylation levels arises from technique differences rather than conditions (see Figure 6). Perturbation samples (e.g. writer knockdown) were excluded at this step to retain only wild-type samples, preserving a consistent basal m^6^A pattern after integration. The site calling model was then fitted to these combined samples, and m^6^A site probabilities were calculated for IDR analysis. The rationale for using m^6^A site probability over p-value is detailed in Supplementary Figure 1. Only sites with total coverage over 30 in both samples were included in the IDR analysis to reduce noise from gene expression variability. For condition-specific m^6^A sites, an alternative integration strategy was provided by summing while separating different conditions and tissues. These condition-specific results can be downloaded at m6AConquer web server.

### Empirical Evaluation of m6A Site Calling and Integration Strategies

Multiple benchmarks were designed to empirically justify the selection of site-calling and integration strategies. For site calling, in addition to the BIC approach defined above, the average distance of called sites to the nearest splicing junction was calculated to evaluate the performance of site-calling methods (Figures 4c and 4d). To evaluate the quality of different m^6^A profiling techniques in terms of site prediction accuracy, AUROC (Area Under the Receiver Operating Characteristic curve) was calculated using the posterior prediction of m^6^A site probabilities inferred from BBmix on combined samples, where the ground truth was defined by the high-confidence integrated sites. For benchmarking alternative integration methods and databases, eTAM-seq and GLORI were used as ground truth due to their high fidelity.

**Figure 4.**
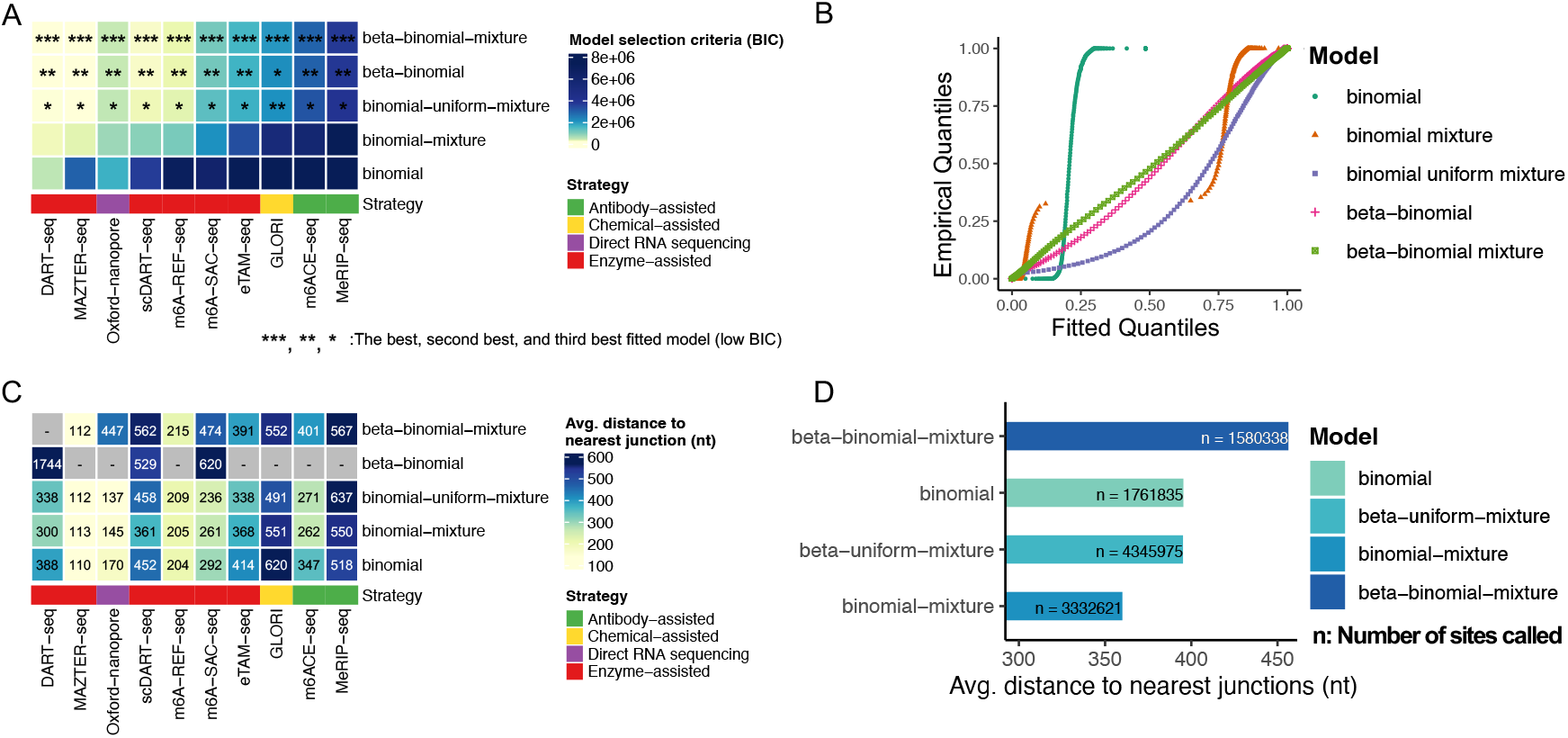
Evaluation of Statistical Models for m6A Site Calling (**A**) Heatmap comparing the empirical fit for various statistical models across m6A detection techniques, using the Bayesian Information Criterion (BIC). Lower BIC indicates a better fit, with the beta-binomial mixture model consistently performing best (***, **, * representing the top three models). (**B**) Quantile-Quantile (QQ) plots comparing empirical versus fitted quantiles for m6A methylation ratios on m6A-REF-seq, demonstrating the models’ ability to capture m6A level distribution. (**C**) Heatmap showing the average distance of called m^6^A sites to the nearest splicing junction across different models and techniques. “-” indicates no sites called. (**D**) Bar chart comparing the average distance to the nearest splicing junction for m6A sites across models in (C). The beta-binomial mixture model identifies a comparable number of sites with binomial, while with the greatest average distance (450 nt) to splicing junctions, indicating its effectiveness in identifying biologically relevant m^6^A sites.

To further assess the reliability of biological interpretations using the high-confidence integrated sites, genomic feature importance from random forest (RF) classification models was calculated. We chose RF as the machine learning algorithm to rank genomic features because of its strength in capturing non-linear correlations and interactions in high-dimensional feature space. The RF classifiers were built using the high-confidence integrated sites (IDR < 0.05) derived from four pairs of independent techniques (Figure 7A) but without p-value filtering. These datasets were selected due to their reproducible regions spanned both the positive and negative parts for both techniques. The negative data in the ML classification were defined by an m^6^A site probability < 0.5 in both techniques. AUROC of RF classifiers was computed using ten-fold cross-validation to evaluate the prediction performance. Feature importance in Random Forest models was assessed by measuring the reduction in Gini impurity when a specific feature is removed, as implemented in the *randomForest* R package.

The predictive genomic features used in the RF models were features of intron-exon architectures extracted using the ’genomeDerivedFeatures’ function in the *predictiveFeatures* R package (55) with input annotation from hg38 txdb. The extracted features included a comprehensive combination of different transcriptomic regions (e.g., exon, intron, CDS, 5’UTR, and 3’UTR) and their properties (length, relative positions, distance to the five-prime and three-prime ends) (see Figure 7C). Additionally, a new feature was introduced to represent the distance to the nearest splicing junction, which has been suggested as a driving factor in m^6^A formation (9-11).

## RESULTS & DISCUSSION

### Comparative Analysis of m^6^A Profiling Techniques

The Pearson correlation between methylation levels (defined by the beta value or the ratio of m^6^A counts to total coverage) was calculated across all 2,788,705 sites in human samples, comparing combined samples from each technique (Figure 3A). Correlations were computed by considering only pairs of observations where both sites had a total coverage greater than 30, excluding any positions with missing data. Overall, GLORI and eTAM-seq demonstrated the highest average correlation with other techniques, whereas DART-seq and scDART-seq showed the lowest. This finding supports the current understanding that eTAM-seq and GLORI, as absolute quantitative methods, are more accurate. In contrast, DART-seq and scDART-seq, which rely heavily on the YTH domain of m^6^A reader proteins, may not capture a representative global m^6^A profile as effectively as other methods.

The numbers of integrated high-confidence sites between independent techniques were then examined with an upset plot (Figure 3D). The integrated outcomes are defined with an IDR threshold of < 0.05, together with the adjusted p-value < 0.05 in both techniques. The input signal for IDR calculation is m^6^A site probability inferred from the combined samples. Only pairs between independent techniques are displayed to avoid counting overlaps due to technical confounding factors. The upset plot displays the top six techniques with the highest number of reproduced sites. The greatest number of cross-technique intersected sets belonged to eTAM-seq and GLORI. The second largest set belongs to the intersection between eTAM-seq, GLORI, and m6ACE-seq. m6A-SAC-seq and MeRIP-seq also exhibit a moderate number of overlapped sites with other techniques. Techniques not displayed in the graph generally showed weaker alignment with other techniques in terms of the IDR integration outcomes.

Next, the site prediction accuracy of each m^6^A profiling technique was assessed using Bayesian classification based on the fitted BBmix models. Bayes classifiers were trained on the combined samples for each m^6^A detection method, with the predictive values being the m^6^A site probabilities. The evaluation used high-confidence integrated sites as the ground truth for positive sites, while all other sites in the human reference were considered negative. We found that Bayesian classification performed best for eTAM-seq (AUC = 0.99) and GLORI (AUC = 0.97), while DART-seq (AUC = 0.55) and scDART-seq (AUC = 0.48) showed lower performance (Figure 3B). Notably, MeRIP-seq’s ranking improved compared to the m^6^A level correlation benchmark, likely due to its high coverage, which suggests that its non-absolute nature is more effective for site identification than for quantification.

### Evaluation and Selection of Optimal Site Calling Methods

To empirically justify the choice of the BBmix model as the uniformed solution for site calling and normalization across all m^6^A quantification techniques, two types of benchmarks were conducted to compare it against alternative statistical models. First, model empirical fit is evaluated using the Bayesian Information Criterion (BIC) and Quantile-Quantile (QQ) plots. The results showed that the BBmix model consistently dominated all other models across nearly every sample from all detection techniques (Figure 4A and 4B; Supplementary Figure 3). This outcome is not surprising, as BBmix effectively captures the over-dispersion of the binomial model, significantly enhancing the model’s expressiveness.

Subsequently, the evaluation of the biological significance of the sites called using different methods is performed, with all methods calling sites under the threshold of adjusted p-value less than 0.05. The average distance of the positively identified m^6^A sites to the nearest splicing junction was used as a measure of the scientific significance. The result demonstrates that, across most techniques, BBmix returned a comparable number of positive sites to competing methods while yielding the highest average distance to splicing junctions among the majority of techniques (Figure 4C and 4D). Additionally, for GLORI and eTAM-seq, BBmix called significantly more sites than the binomial method used in their original pipelines (Supplementary Figure 1A). Moreover, the topology of new sites identified only by BBmix in GLORI and eTAM-seq aligned well with the known topology of m6A (Supplementary Figure 1B). Therefore, this observation suggests that the m6A site-calling threshold defined by BBmix is generally robust across techniques and often better represents the underlying biological signals.

### Evaluation of Integration Approaches for m^6^A Profiling Techniques

An additional empirical benchmark was conducted to compare methods of data integration across different m^6^A profiling techniques. The ten m^6^A detection techniques were grouped into four categories based on their experimental mechanisms: antibody-assisted, enzyme-assisted, direct-RNA sequencing, and chemical-assisted (Figure 3C). The IDR approach was used to integrate data, comparing results between orthogonal techniques (across different categories) and dependent techniques (within the same category), with reproducible sites defined as having an IDR < 0.05. The results showed that both orthogonal and dependent integrations could yield significant, and in some cases, globally positive results (e.g., GLORI vs. eTAM-seq for orthogonal integration and MeRIP-seq vs. m6ACE-seq for dependent integration) (Figures 4a). However, the integration of orthogonal techniques consistently outperformed that of dependent techniques, as indicated by the higher predictive performance for high-quality sites in GLORI and eTAM-seq data (Figure 5B). IDR plots between all technique pairs were put in Supplementary Figure 2.

**Figure 5.**
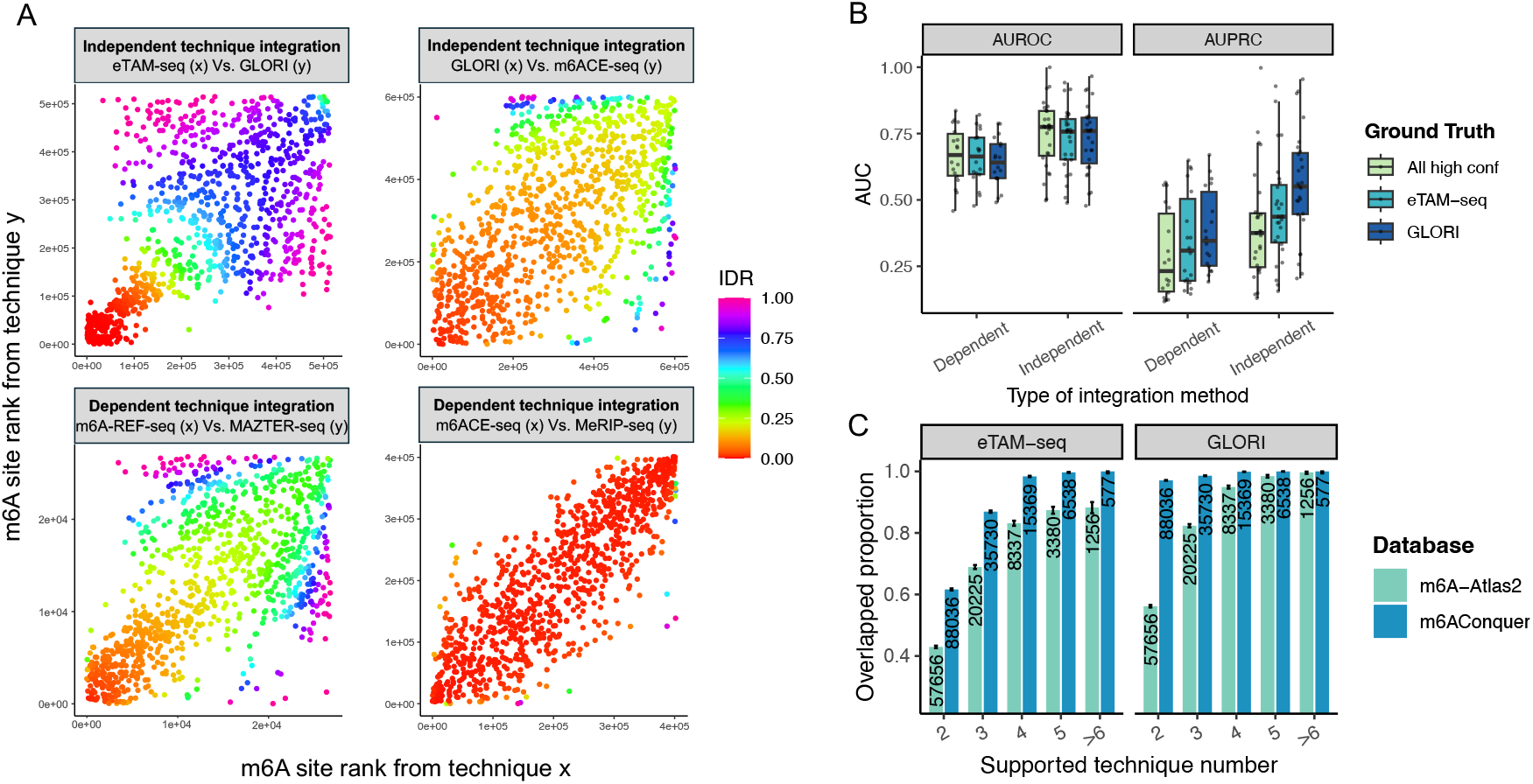
Evaluation of Independent and Dependent Methods for Cross Platform m^6^A Integration (**A**) Scatter plots showing m^6^A site rank integration between independent (top) and dependent (bottom) technique pairs, with sites color-coded by Irreproducible Discovery Rate (IDR). The x and y axes represent the rank of m^6^A site probability calculated from the respective detection technique. (**B**) Box plots comparing AUROC and AUPRC for m^6^A site predictions using IDR from independent versus dependent integration methods. Independent integration consistently outperforms dependent methods across three ground truths (all high-confidence integrated sites, eTAM-seq, and GLORI). (**C**) Bar chart comparing the overlap proportion of m^6^A sites between m6AConquer and m6A-Atlas2 databases across different numbers of supporting techniques. m6AConquer shows higher overlap proportions with GLORI and eTAM-seq, meanwhile have more sites reported at each threshold. Numbers on the bars indicate the count of high-confidence sites supported at the threshold.

Three different input quantities for IDR integration (used in the rank calculation) were benchmarked: m^6^A level (the ratio of m^6^A to total coverage), m^6^A site probability, and -log adjusted p-values. The results indicated that high-confidence integrated sites at IDR < 0.05 were largely shared across all three approaches, with m^6^A site probability and m6A level identifying additional number of high-confidence sites (Supplementary Figure 1C). The unique sites identified by both methods exhibited a reasonable transcript topology distribution (Supplementary Figure 1D). Based on these observations, m^6^A site probability and m6A level are considered the better choices for the input of the IDR integration process. As a result, m^6^A site probability was selected as the default integration method, as it fully accounts for the stochastic nature of the marginal data distribution.

Subsequently, the high-confidence integrated sites in m6AConquer were evaluated in comparison to the human base resolution sites in m6A-Atlas2, one of the popular m^6^A modification database (65). m6A-Atlas2 uses the intersection method (finding overlaps) for cross-technique integration. In contrast, our m6AConquer employs the model-based method through IDR while considers technical orthogonality. The precision of integrated sites in both databases was assessed using high-quality data from GLORI and eTAM-seq. Different thresholds are set based on the number of supporting techniques reported by each database, ranging from 2 to 6. The findings revealed that at each supported level, m6AConquer demonstrated higher precision than m6A-Atlas2 meanwhile reported more sites at each level, except in the category where the support number is *≥6* (Figure 5C). This result suggests that m6AConquer effectively captures a higher number of high-confidence m^6^A sites with greater credibility compared to popular databases that use non-quantitative, intersection-based integration methods.

### Comparison of Biological and Technical Variation in m6A Methylation Levels

In m6AConquer, quantitative m^6^A data were consistently collected across different techniques in a matrix format, thus enabling accurate quantification of variance associated with techniques, conditions, and batches. A conceptually similar analysis was conducted via the mixed effect linear model in the CHIP-Seq data within the ENCODE dataset (62). To intuitively compare sample variance estimates of sites’ m6A levels, the variance was calculated both across different conditions within the same technique and across different techniques within the same condition. Sites with a total coverage less than 30 were again omitted in variance calculation due to the high estimation error of m6A level. When comparing the pooled variances between the two groups, it was found that the variance between techniques within the same condition is, on average, much larger than the variance between conditions within the same technique (Figure 6).

**Figure 6.**
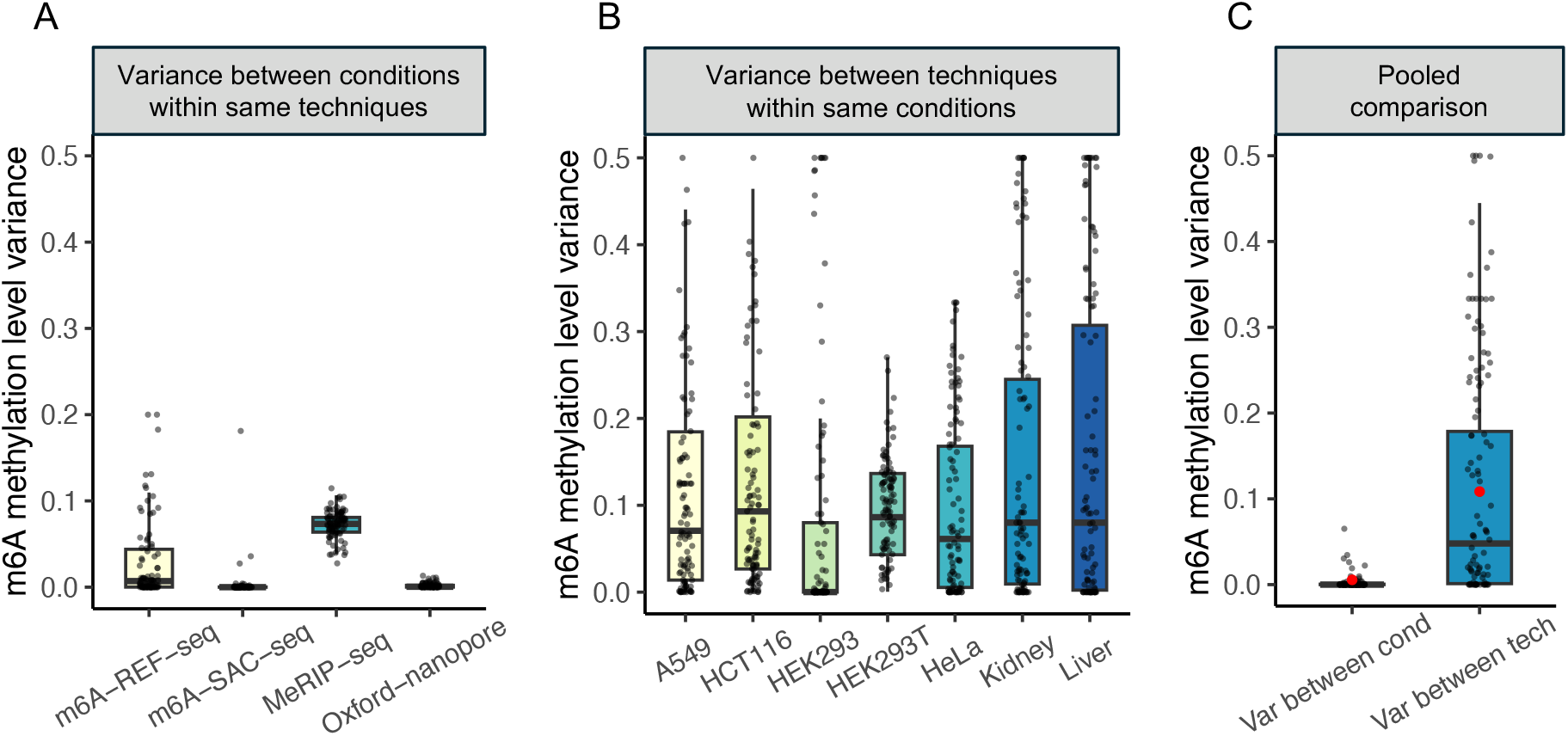
Comparison of m^6^A Methylation Level Variance Across Techniques and Conditions. (**A**) Box plots for variance in m^6^A methylation levels across different conditions within the same technique. (**B**) Box plots for variance across different techniques within the same condition. Conditions without matched multiple techniques are not displayed. (**C**) Pooled comparison showing that variance is significantly higher between techniques (blue) than between conditions (red). This observation suggests that technical differences are a major source of variability in quantitative m^6^A data, demonstrating the importance of integrating data from orthogonal m^6^A detection methods.

This observation supports the current understanding that m^6^A is constitutively installed across all cell lines, with changes in m^6^A levels primarily driven by the intron-exon architecture of expressed transcript isoforms (3,9-11,96). Therefore, m^6^A levels are not expected to vary significantly across most peripheral tissues and cell-line conditions within a species (97). In contrast, the technical variances introduced by different m^6^A detection techniques are relatively larger, due to substantial differences in their experimental instruments. These results also justify the use of combined samples in many analyses presented in this manuscript, since the majority of sources of data variances are still preserved after pooling replicates within techniques.

### Enhanced Biological Interpretability Through Integration of m^6^A Profiling Techniques

By integrating orthogonal m^6^A profiling techniques, a higher level of biological interpretability can be achieved. To demonstrate biological significance, the genomic feature importance ranked by Random Forest Classifiers was calculated based on the high-confidence integrated sites across orthogonal technical pairs. Genomic features are defined by the intron-exon architecture of various combinations of transcriptomic regions (e.g. exons, CDS) and their properties (e.g. length, distance to ends) (see Figure 7C). The resulting feature importance rankings were found to be consistent across all four pairs of orthogonal integration outcomes, with Spearman correlations exceeding 0.9553 (Figure 7A). This result highlights the effective removal of technical artifacts using the IDR integration framework between independent techniques.

**Figure 7.**
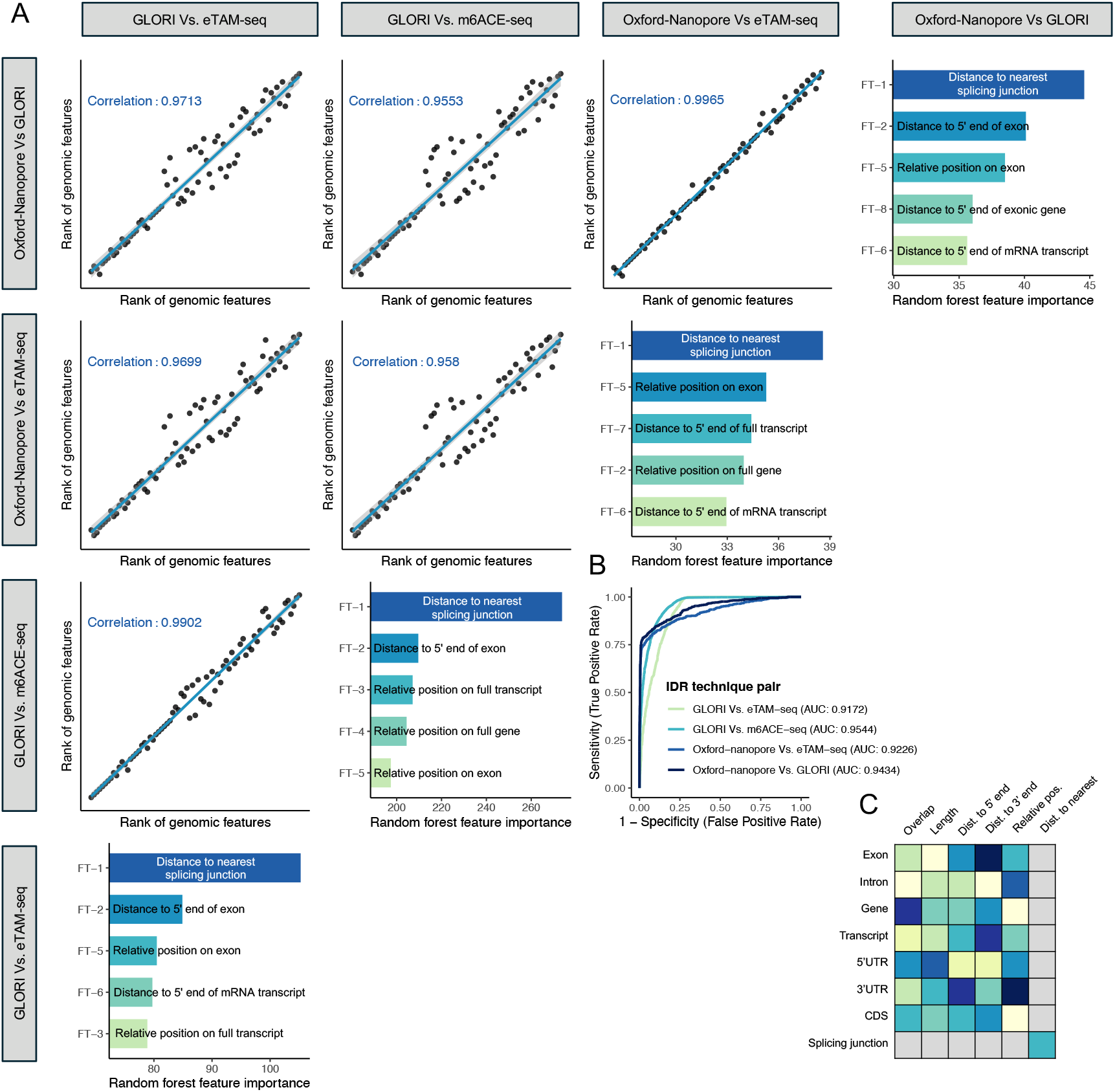
Correlation of Genomic Feature Importance Across High-Confidence Integrated Sites (**A**) Scatter plots showing high correlations in ranks of genomic feature importance between integration results from independent pairs of m^6^A detection techniques. The bar plots list the top 5 important features. All four datasets consistently rank ‘Distance to Nearest Splicing Junction’ as the most important feature in Random Forest models. (**B**) ROC curves for integrated m^6^A site predictions using genomic features, with GLORI vs. m6ACE-seq showing the highest AUC (0.9544), demonstrating strong predictive performance of the models. (**C**) An illustration of the genomic feature representation used. Transcriptomic regions (rows in the heatmap) and properties (columns in the heatmap) are combined to model intron-exon architecture. An additional feature, ‘Distance to Nearest Splicing Junction’, is included, as shown in the lower left corner.

Additionally, it was consistently observed that the most important feature across all four examined pairs is the distance to the nearest splicing junction (see diagonal in Figure 7A). This finding aligns with previous experimental studies, which suggest that m^6^A is predominantly installed through an inhibition mechanism at the splicing junction complex (3,9-11). The spline logistic regression curves were fitted to identify the decision boundary for the single feature. The result indicates the sites with a distance greater than 60 nucleotides from the nearest splicing junction are more likely to be m^6^A sites, and this boundary is consistently supported across all four datasets (see Supplementary Figure 5).

It is worth noting that, other genomic features previously linked to m^6^A distribution, such as exon length, stop codon, and last exon, are full captured by the region property features in the current RF models. However, their lower feature importance suggests they are likely confounding factors related to the distance to the nearest splicing junction rather than primary drivers. Using genomic features and Random Forest alone yields highly accurate m^6^A classification models, with AUROC values between 0.9172 and 0.9544 (Figure 7B). Additionally, Gene Ontology (GO) analysis of high-confidence m^6^A sites from different orthogonal technique pairs (IDR < 0.05, padj < 0.05) revealed consistent enrichment across all three categories, with the most significant terms being DNA replication (biological process), ubiquitin ligase complex (cellular component), and catalytic activity on DNA (molecular function) (Supplementary Figure 4).

## CONCLUSION

To conclude, we present m6AConquer, a database designed for consistent quantification of external m^6^A RNA modification data. This work aims at addressing several key gaps in the field:

1. The absence of a standardized multi-omics data framework to enable coherent data sharing and integration across different platforms, studies, and omics modalities in m^6^A epitranscriptomics research.
2. The missing of a model-based integration framework that acknowledges the (in)dependence of techniques across various m^6^A detection platforms.
3. The omission of a consistent application of statistical methods that account for over-dispersion in site calling and normalization processes.

This manuscript demonstrates the significance of incorporating consistent data framework and robust statistical techniques into the epirtanscriptomic data resources through evidence-based benchmarks. The proposed model-based integration method identified a significant number of reproducible sites across various technical pairs. Among them, integration between independent technical categories are showed to be more accurate than between dependent categories. Additionally, the uniform site-calling and normalization approach, which accounts for over-dispersion in m^6^A levels, was shown to deliver potentially superior performance than popular method across ten m^6^A profiling techniques. Furthermore, the model-based integration using IDR was shown to be potentially more reliable than intersection-based integration through quantification of statistical uncertainties, as evidenced by the cross-database comparison with m6A-Atlas2. It is worth noting that, GLORI and eTAM-seq emerged to be the most accurate m^6^A profiling techniques in our evaluation, consistently dominating other detection methods in terms of prediction accuracy for high-confidence m^6^A sites.

We further conducted an intuitive analysis of our integrated m^6^A sites to rank genomic factors within intron-exon architecture based on their predictive importance in machine learning models for m^6^A sites. This analysis consistently identified the distance to the nearest intron-exon junction (IEJ) as the most important factor in predicting m^6^A sites across all examined datasets of reproducible sites from orthogonal techniques. This finding suggests that the distance to the IEJ is likely the primary genomic factor driving m6A installation. Other features, such as exon length and relative position within the 5’ UTR, CDS, and 3’ UTR, appear to be confounding factors correlated to the distance to the IEJ. This suggests a need to critically reassess the current practice of using transcript topology—such as that drawn by the popular *guitar* package (98) —as the main omics feature used for explaining m6A data.

In the upcoming upgrade of m6AConquer, additional multi-omics modalities will be introduced beyond the current m6A-Omics and transcriptomics. A key focus will be incorporating genetic variant information into the Multi-Assay Experiment framework, enabling consistent m6A-QTL analysis across different datasets and techniques. Given the significant role of the intron-exon junction (IEJ) in m6A installation, the in-silico mutation approach will be enhanced to better account for this mechanism. Current RNA modification databases use sequence-based machine learning models with short sequence inputs (20-500bp) for functional annotation of m6A sites, but these models often fail to capture the broader effects of changes in the IEJ, as shown by m^6^A-QTL studies (39,99). Consequently, in-silico mutation information has not been included in this version of m6AConquer due to concerns about its consistency with m6A-QTL data and overall reliability. In the future development of m6AConeuer, the potential included functional annotation by ML will be subjected to careful empirical testing and calibration.

In conclusion, the m6AConquer project has established a unifying data resource that enables consistent quantification and systematic integration of 10 quantitative m^6^A detection techniques. High-quality, user-friendly data structures are assembled to facilitate omics-based analysis of m^6^A RNA modification. This resource is designed to support both computational and experimental researchers in epitranscriptomics by providing clean, comprehensive data, along with empirically validated normalization and integration procedures to minimize artifacts. Deriving biological conclusions supported by multiple independent techniques is crucial, and m6AConquer is dedicated to providing summarized raw information and automated integration outcomes to achieve this goal. The database, along with detailed instructional tutorials, is available at http://rnamd.org/m6aconquer.

## AUTHOR CONTRIBUTIONS

Xichen Zhao: Data Collection & Processing, Formal analysis, Validation, Visualization, Presentation, Methodology, Writing—review & editing. Haokai Ye: Website Design & Development. Daniel Rigden: Writing—review & editing. Tenglong Li: Writing—review & editing. Zhen Wei: Conceptualization, Methodology, Supervision, Writing—original draft.

## DATA AVAILABILITY

The m6AConquer database is freely accessible at http://rnamd.org/m6aconquer. The OmixM6A R package, which facilitates the reproduction of site calling and normalization methods or the fitting of alternative models, is available at GitHub (https://github.com/ZW-xjtlu/OmixM6A) and archived at Zenodo (DOI:10.5281/zenodo.13337113). The source code for processing raw data in m6AConquer is provided at GitHub (https://github.com/XichenZhao0223/m6AConquer) with an accompanying Zenodo archive (DOI: 10.5281/zenodo.13337723). Additional supporting data are available in the Supplementary Material.

## ACKNOWLEDGEMENT

We extend our gratitude to Professor Jia Meng and Dr. Francesco Zonta for providing critical feedbacks. We also thank Shuhe Liu for her valuable suggestions on the manuscript presentation. We thank Dr Bowen Song for providing us with the processed scDART-seq data.

