## Supplementary Material for "m6AConquer: a Data Resource for Unified Quantification and Integration of m6A Detection Techniques"

### Supplementary Figures

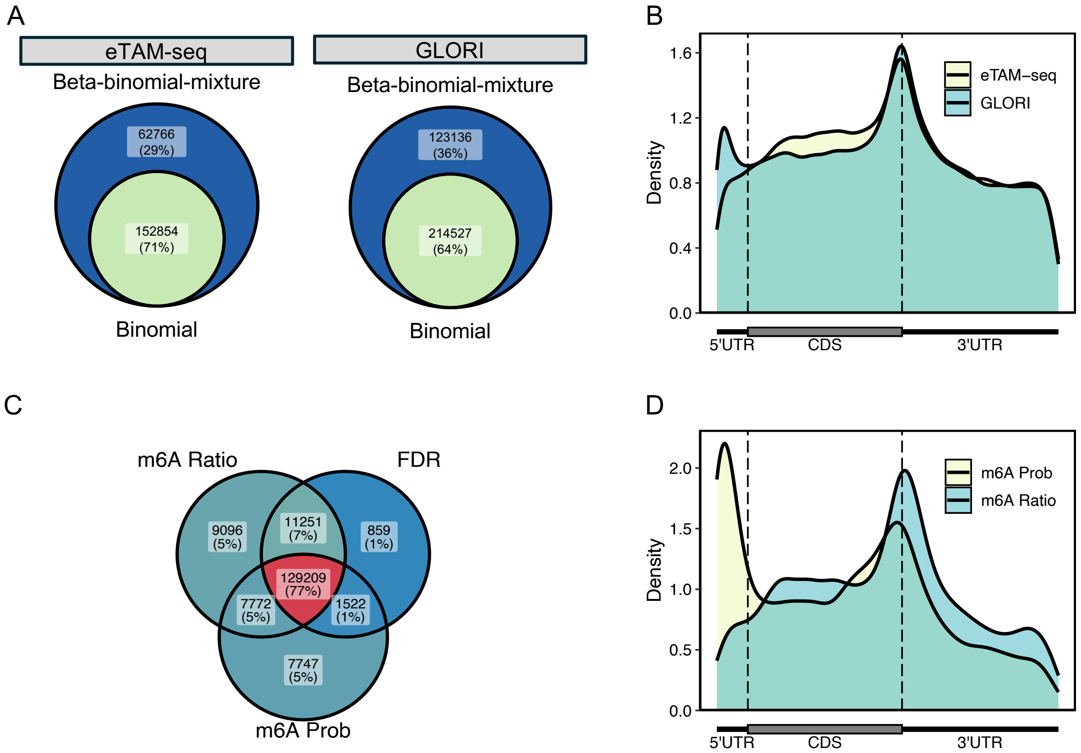


**Figure 1.** Evaluation of the site calling methods and IDR integration inputs.

**(A)** Venn diagram comparing m6A site-calling methods in eTAM-seq and GLORI data. The outer circles represent sites identified by binomial mixture models (blue), and the overlapped section in the middle represents sites identified by both binomial and beta-binomial mixture models (green), with percentages indicating the proportion of each subset.

**(B)** Metagene topology plot indicating the distribution of m6A sites across gene regions exclusively identified by beta-binomial mixture models from the Venn diagram in A. The X-axis represents gene regions: 5'UTR (untranslated region), CDS (coding sequence), and 3'UTR. The Y-axis shows the site density.

**(C)** Venn diagram showing the overlap of reproducible m6A sites identified using three different inputs for IDR (irreproducible discovery rate) calculation - m6A Ratio, m6A Prob ($m^6A$ site probabilities), and FDR. The central overlapped section shows sites identified by all three metrics, with percentages indicating the proportion of each subset.

**(D)** Metagene topology plot highlighting the distributions of m6A sites identified exclusively through m6A ratios and $m^6A$ site probabilities as indicated in the Venn diagram in C. The X-axis and Y-axis are the same as in B.

Note: The percentages inside the circles in Venn diagrams A and C indicate the proportion of sites relative to the total number of sites identified by the corresponding or combination of methods.

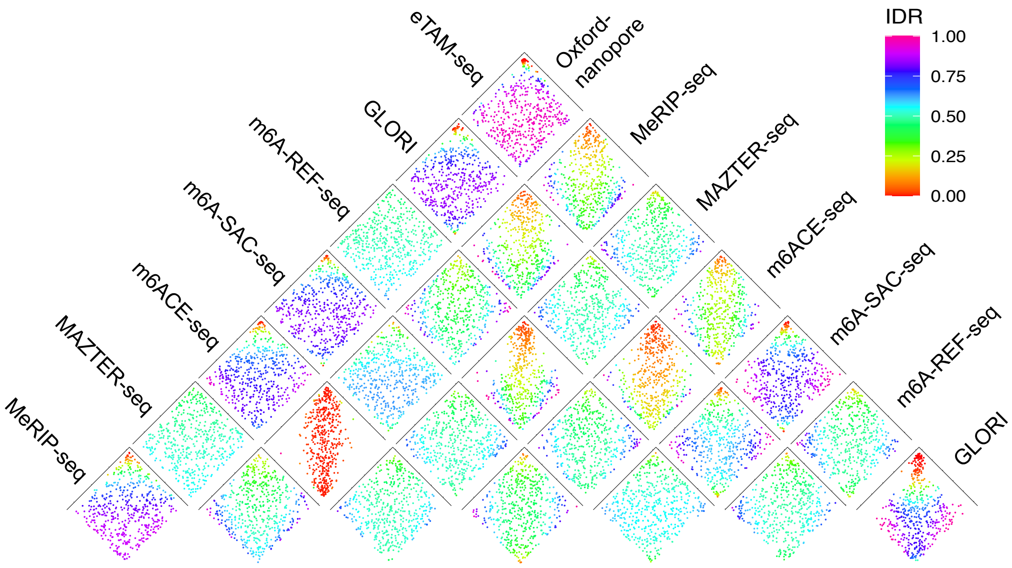


**Figure 2.** Scatter plots showing ranks of m6A probabilities between all integrated technique pairs, with sites colour-coded by Irreproducible Discovery Rate (IDR).

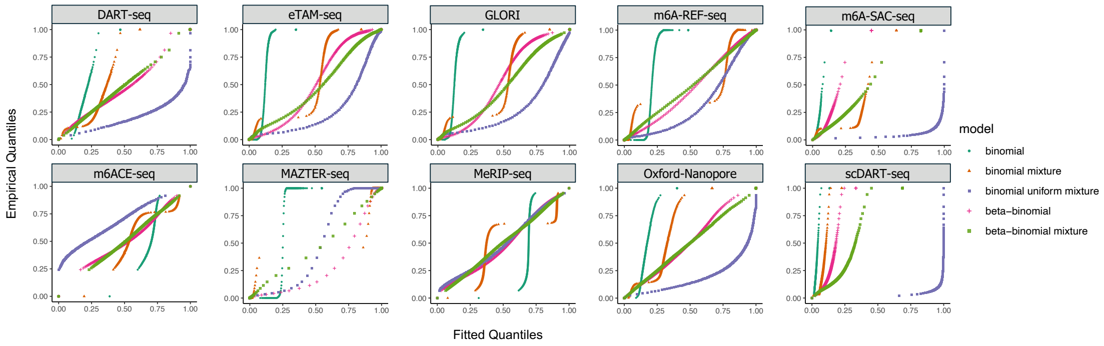


**Figure 3.** Goodness of fit for all candidate models. The x-axis of each plot represents the Fitted Quantiles of m6A ratios, and the y-axis represents the Empirical Quantiles of m6A ratios. The closer the points of a model follow the 45-degree line, the better the model fits the data from that sequencing technique. Beta-binomial mixture models are closest to the 45-degree line in all techniques among all candidate models.

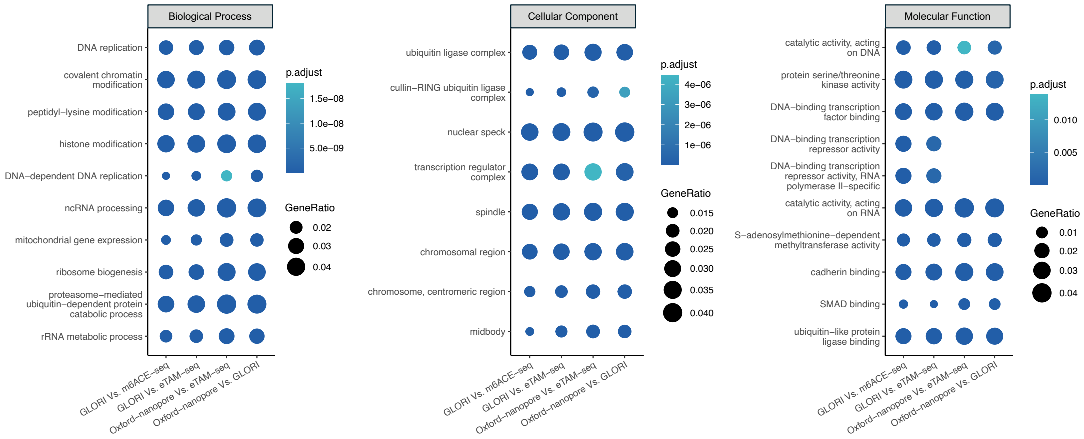


**Figure 4.** Gene ontology (GO) enrichment analysis of integrated high-confidence sites. The dot plots visualize the GO enrichment for genes with reproducible m6A sites identified through IDR integration between 4 pairs of independent techniques (GLORI and m6ACE-seq, GLORI and eTAM-seq, Oxford-nanopore sequencing and eTAM-seq, and Oxford-nanopore sequencing and GLORI). Dot color represents the adjusted p-value (p.adjust) with darker shades indicating more significant enrichment. Dot size indicates the ratio of genes within a particular GO term to the total number of enriched genes (GeneRatio).

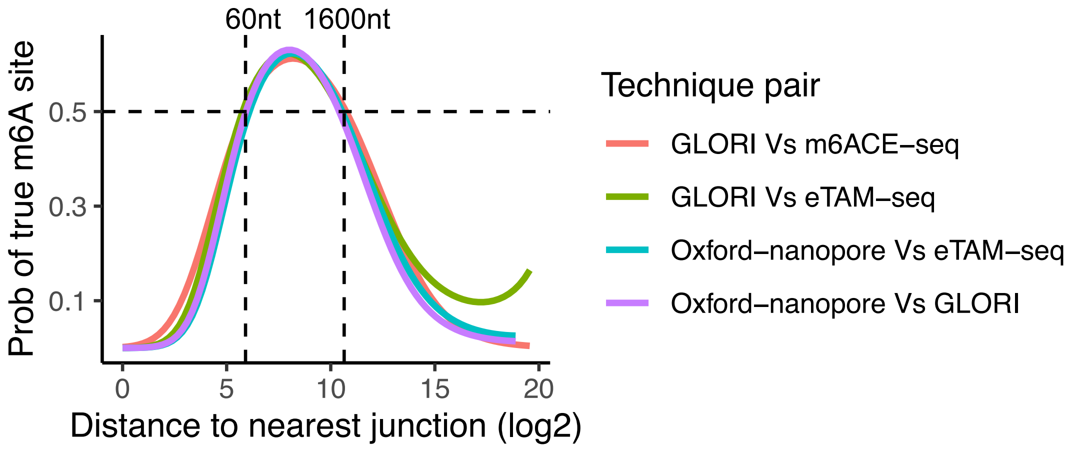


**Figure 5.** Spline logistic regression analysis of reproducible m6A sites relative to the distances to the nearest splicing junction. The curves demonstrate the probability of identifying reproducible m6A sites against the log2 scale distances to the nearest splicing junction. Each curve represents the reproducible m6A sites from one IDR integration technique pairs (GLORI and m6ACE-seq, GLORI and eTAM-seq, Oxford-nanopore sequencing and eTAM-seq, and Oxford-nanopore sequencing and GLORI). The horizontal dashed line marks the 0.5 probability threshold. The vertical dashed lines represent the decision boundaries of 60nt and 1600nt.

### Supplementary Table

The supplementary table contains the information on data used in the analysis, except for scDART-seq. scDART-seq are retrieved from GSE180954 datasets, including 1000 sequencing data from SRR15268425 to SRR15270105.

| Run Accession | Sample Accession | Experiment Accession | Type | Technique | Condition |
|---|---|---|---|---|---|
| SRR15268877 | GSM5479237 | GSE180954 | Wild type | DART-seq | HEK293T |
| SRR15268878 | GSM5479238 | GSE180954 | Wild type | DART-seq | HEK293T |
| SRR15268879 | GSM5479239 | GSE180954 | Wild type | DART-seq | HEK293T |
| SRR9940470 | GSM4024125 | GSE125780 | Wild type | DART-seq | HEK293T |
| SRR9940471 | GSM4024126 | GSE125780 | Wild type | DART-seq | HEK293T |
| SRR9940472 | GSM4024127 | GSE125780 | Wild type | DART-seq | HEK293T |
| SRR9940473 | GSM4024128 | GSE125780 | Wild type | DART-seq | HEK293T |
| SRR21070395 | GSM6463607 | GSE201064 | IVT | eTAM-seq | HeLa |
| SRR21070396 | GSM6463606 | GSE201064 | IVT | eTAM-seq | HeLa |
| SRR21070397 | GSM6463605 | GSE201064 | IVT | eTAM-seq | HeLa |
| SRR21070398 | GSM6463605 | GSE201064 | IVT | eTAM-seq | HeLa |
| SRR21070399 | GSM6463604 | GSE201064 | Perturbation | eTAM-seq | HeLa |
| SRR21070400 | GSM6463603 | GSE201064 | Perturbation | eTAM-seq | HeLa |
| SRR21070401 | GSM6463602 | GSE201064 | Perturbation | eTAM-seq | HeLa |
| SRR21070402 | GSM6463602 | GSE201064 | Perturbation | eTAM-seq | HeLa |
| SRR21070403 | GSM6463601 | GSE201064 | Wild type | eTAM-seq | HeLa |
| SRR21070404 | GSM6463600 | GSE201064 | Wild type | eTAM-seq | HeLa |
| SRR21070405 | GSM6463599 | GSE201064 | Wild type | eTAM-seq | HeLa |
| SRR21070406 | GSM6463599 | GSE201064 | Wild type | eTAM-seq | HeLa |
| SRR21356198 | GSM6432643 | GSE210563 | Wild type | GLORI | HEK293T |
| SRR21356199 | GSM6432642 | GSE210563 | Wild type | GLORI | HEK293T |
| SRR21356201 | GSM6432640 | GSE210563 | IVT | GLORI | HEK293T |
| SRR21356202 | GSM6432641 | GSE210563 | IVT | GLORI | HEK293T |
| SRR21356250 | GSM6432591 | GSE210563 | Wild type | GLORI | HEK293T |
| SRR21356251 | GSM6432590 | GSE210563 | Wild type | GLORI | HEK293T |
| SRR11015605 | GSM4294990 | GSE124509 | Perturbation | m6ACE-seq | HEK293T |
| SRR11015606 | GSM4294991 | GSE124509 | Perturbation | m6ACE-seq | HEK293T |
| SRR11015607 | GSM4294992 | GSE124509 | Perturbation | m6ACE-seq | HEK293T |
| SRR11015608 | GSM4294993 | GSE124509 | Perturbation | m6ACE-seq | HEK293T |
| SRR11015609 | GSM4294994 | GSE124509 | Perturbation | m6ACE-seq | HEK293T |
| SRR11015610 | GSM4294995 | GSE124509 | Perturbation | m6ACE-seq | HEK293T |
| SRR11015611 | GSM4294996 | GSE124509 | Perturbation | m6ACE-seq | HEK293T |
| SRR11015612 | GSM4294997 | GSE124509 | Perturbation | m6ACE-seq | HEK293T |
| SRR11015613 | GSM4294998 | GSE124509 | Perturbation | m6ACE-seq | HEK293T |
| SRR8380776 | GSM3535501 | GSE124509 | Wild type | m6ACE-seq | HEK293T |
| SRR8380778 | GSM3535503 | GSE124509 | Wild type | m6ACE-seq | HEK293T |
| SRR8380780 | GSM3535505 | GSE124509 | Wild type | m6ACE-seq | HEK293T |

| | | | | | |
|---|---|---|---|---|---|
| SRR8380782 | GSM3535507 | GSE124509 | Perturbation | m6ACE-seq | HEK293T |
| SRR8380784 | GSM3535509 | GSE124509 | Perturbation | m6ACE-seq | HEK293T |
| SRR8380786 | GSM3535511 | GSE124509 | Perturbation | m6ACE-seq | HEK293T |
| SRR8380788 | GSM3535513 | GSE124509 | Perturbation | m6ACE-seq | HEK293T |
| SRR8380800 | GSM3535525 | GSE124509 | Perturbation | m6ACE-seq | HEK293T |
| SRR8380802 | GSM3535527 | GSE124509 | Perturbation | m6ACE-seq | HEK293T |
| SRR8380804 | GSM3535529 | GSE124509 | Perturbation | m6ACE-seq | HEK293T |
| SRR8380806 | GSM3535531 | GSE124509 | Perturbation | m6ACE-seq | HEK293T |
| SRR8380808 | GSM3535533 | GSE124509 | Perturbation | m6ACE-seq | HEK293T |
| SRR8380810 | GSM3535535 | GSE124509 | Perturbation | m6ACE-seq | HEK293T |
| SRR9060600 | GSM3768322 | GSE124509 | Perturbation | m6ACE-seq | HEK293T |
| SRR9060602 | GSM3768324 | GSE124509 | Perturbation | m6ACE-seq | HEK293T |
| SRR9060604 | GSM3768326 | GSE124509 | Perturbation | m6ACE-seq | HEK293T |
| SRR9060606 | GSM3768328 | GSE124509 | Perturbation | m6ACE-seq | HEK293T |
| SRR9060608 | GSM3768330 | GSE124509 | Perturbation | m6ACE-seq | HEK293T |
| SRR9060610 | GSM3768332 | GSE124509 | Perturbation | m6ACE-seq | HEK293T |
| SRR9060612 | GSM3768334 | GSE124509 | Perturbation | m6ACE-seq | HEK293T |
| SRR9060614 | GSM3768336 | GSE124509 | Perturbation | m6ACE-seq | HEK293T |
| SRR9060616 | GSM3768338 | GSE124509 | Perturbation | m6ACE-seq | HEK293T |
| SRR9060618 | GSM3768340 | GSE124509 | Perturbation | m6ACE-seq | HEK293T |
| SRR9060620 | GSM3768342 | GSE124509 | Perturbation | m6ACE-seq | HEK293T |
| SRR9060622 | GSM3768344 | GSE124509 | Perturbation | m6ACE-seq | HEK293T |
| SRR9060624 | GSM3768346 | GSE124509 | Perturbation | m6ACE-seq | HEK293T |
| SRR9060626 | GSM3768348 | GSE124509 | Perturbation | m6ACE-seq | HEK293T |
| SRR9060628 | GSM3768350 | GSE124509 | Perturbation | m6ACE-seq | HEK293T |
| SRR9060630 | GSM3768352 | GSE124509 | Perturbation | m6ACE-seq | HEK293T |
| SRR9060632 | GSM3768354 | GSE124509 | Perturbation | m6ACE-seq | HEK293T |
| SRR9060634 | GSM3768356 | GSE124509 | Perturbation | m6ACE-seq | HEK293T |
| SRR9060636 | GSM3768358 | GSE124509 | Perturbation | m6ACE-seq | HEK293T |
| SRR9060638 | GSM3768360 | GSE124509 | Perturbation | m6ACE-seq | HEK293T |
| SRR9060640 | GSM3768362 | GSE124509 | Perturbation | m6ACE-seq | HEK293T |
| SRR9073880 | GSM3535515 | GSE124509 | Perturbation | m6ACE-seq | HEK293T |
| SRR9073882 | GSM3535517 | GSE124509 | Perturbation | m6ACE-seq | HEK293T |
| SRR9905081 | GSM4007645 | GSE124509 | Perturbation | m6ACE-seq | HEK293T |
| SRR9905083 | GSM4007647 | GSE124509 | Perturbation | m6ACE-seq | HEK293T |
| SRR9905085 | GSM4007649 | GSE124509 | Perturbation | m6ACE-seq | HEK293T |
| SRR9905087 | GSM4007651 | GSE124509 | Perturbation | m6ACE-seq | HEK293T |
| SRR9905089 | GSM4007653 | GSE124509 | Perturbation | m6ACE-seq | HEK293T |
| SRR9905091 | GSM4007655 | GSE124509 | Perturbation | m6ACE-seq | HEK293T |
| SRR14765592 | GSM5365409 | GSE151028 | Wild type | m6A-REF-seq | HEK293T |
| SRR14765594 | GSM5365411 | GSE151028 | Wild type | m6A-REF-seq | HEK293T |
| SRR14765614 | GSM5365431 | GSE151028 | Wild type | m6A-REF-seq | HEK293T |
| SRR14765615 | GSM5365432 | GSE151028 | Wild type | m6A-REF-seq | HEK293T |
| SRR14765616 | GSM5365433 | GSE151028 | Wild type | m6A-REF-seq | HEK293T |

| | | | | | |
|---|---|---|---|---|---|
| SRR8450805 | GSM3566973 | GSE125240 | Wild type | m6A-REF-seq | HEK293 |
| SRR8450807 | GSM3566975 | GSE125240 | Wild type | m6A-REF-seq | HEK293 |
| SRR8450809 | GSM3566977 | GSE125240 | Wild type | m6A-REF-seq | HEK293 |
| SRR8450811 | GSM3566979 | GSE125240 | Wild type | m6A-REF-seq | Brain |
| SRR8450813 | GSM3566981 | GSE125240 | Wild type | m6A-REF-seq | Liver |
| SRR8450815 | GSM3566983 | GSE125240 | Wild type | m6A-REF-seq | Kidney |
| SRR13167131 | GSM4949959 | GSE162357 | Wild type | m6A-SAC-seq | HEK293 |
| SRR13167133 | GSM4949961 | GSE162357 | Wild type | m6A-SAC-seq | HEK293 |
| SRR13167136 | GSM4949964 | GSE162357 | Wild type | m6A-SAC-seq | HPSC |
| SRR13167138 | GSM4949966 | GSE162357 | Wild type | m6A-SAC-seq | HPSC |
| SRR13167142 | GSM4949970 | GSE162357 | Wild type | m6A-SAC-seq | HPSC |
| SRR13167144 | GSM4949972 | GSE162357 | Wild type | m6A-SAC-seq | HPSC |
| SRR13167148 | GSM4949976 | GSE162357 | Wild type | m6A-SAC-seq | HPSC |
| SRR13167150 | GSM4949978 | GSE162357 | Wild type | m6A-SAC-seq | HPSC |
| SRR13167154 | GSM4949982 | GSE162357 | Wild type | m6A-SAC-seq | HPSC |
| SRR13167156 | GSM4949984 | GSE162357 | Wild type | m6A-SAC-seq | HPSC |
| SRR13168858 | GSM4950342 | GSE162357 | Wild type | m6A-SAC-seq | HEK293 |
| SRR13168860 | GSM4950344 | GSE162357 | Wild type | m6A-SAC-seq | HEK293 |
| SRR13168864 | GSM4950347 | GSE162357 | Wild type | m6A-SAC-seq | HeLa |
| SRR13168866 | GSM4950349 | GSE162357 | Wild type | m6A-SAC-seq | HeLa |
| SRR13168877 | GSM4950359 | GSE162357 | Wild type | m6A-SAC-seq | HepG2 |
| SRR13168879 | GSM4950361 | GSE162357 | Wild type | m6A-SAC-seq | HepG2 |
| SRR18280446 | GSM5941845 | GSE198246 | Wild type | m6A-SAC-seq | HeLa |
| SRR18280447 | GSM5941844 | GSE198246 | Wild type | m6A-SAC-seq | HeLa |
| SRR18280448 | GSM5941844 | GSE198246 | Wild type | m6A-SAC-seq | HeLa |
| SRR18280453 | GSM5941845 | GSE198246 | Wild type | m6A-SAC-seq | HeLa |
| SRR18280454 | GSM5941841 | GSE198246 | Wild type | m6A-SAC-seq | HeLa |
| SRR18280455 | GSM5941841 | GSE198246 | Wild type | m6A-SAC-seq | HeLa |
| SRR18280456 | GSM5941841 | GSE198246 | Wild type | m6A-SAC-seq | HeLa |
| SRR18280457 | GSM5941840 | GSE198246 | Wild type | m6A-SAC-seq | HeLa |
| SRR18280458 | GSM5941840 | GSE198246 | Wild type | m6A-SAC-seq | HeLa |
| SRR18280459 | GSM5941840 | GSE198246 | Wild type | m6A-SAC-seq | HeLa |
| SRR8903046 | GSM3723216 | GSE122961 | Wild type | MAZTER-seq | hESC |
| SRR8903047 | GSM3723217 | GSE122961 | Wild type | MAZTER-seq | hESC |
| SRR8903048 | GSM3723218 | GSE122961 | Wild type | MAZTER-seq | hESC |
| CRR042278 | CRX037634 | CRA001315 | Wild type | MeRIP-seq | AdrenalGland |
| CRR042280 | CRX037636 | CRA001315 | Wild type | MeRIP-seq | Aorta |
| CRR042282 | CRX037638 | CRA001315 | Wild type | MeRIP-seq | Brainstem |
| CRR042284 | CRX037640 | CRA001315 | Wild type | MeRIP-seq | Cerebellum |
| CRR042286 | CRX037642 | CRA001315 | Wild type | MeRIP-seq | Cerebrum |
| CRR042290 | CRX037646 | CRA001315 | Wild type | MeRIP-seq | Heart |
| CRR042292 | CRX037648 | CRA001315 | Wild type | MeRIP-seq | Hypothalamus |
| CRR042294 | CRX037650 | CRA001315 | Wild type | MeRIP-seq | Liver |
| CRR042296 | CRX037652 | CRA001315 | Wild type | MeRIP-seq | Lung |

| | | | | | |
|---|---|---|---|---|---|
| CRR042298 | CRX037654 | CRA001315 | Wild type | MeRIP-seq | Muscle |
| CRR042300 | CRX037656 | CRA001315 | Wild type | MeRIP-seq | Prostate |
| CRR042302 | CRX037658 | CRA001315 | Wild type | MeRIP-seq | Skin |
| CRR042304 | CRX037660 | CRA001315 | Wild type | MeRIP-seq | Skin |
| CRR042306 | CRX037662 | CRA001315 | Wild type | MeRIP-seq | Stomach |
| CRR042308 | CRX037664 | CRA001315 | Wild type | MeRIP-seq | Stomach |
| CRR042310 | CRX037666 | CRA001315 | Wild type | MeRIP-seq | Testis |
| CRR042312 | CRX037668 | CRA001315 | Wild type | MeRIP-seq | ThyroidGland |
| CRR042314 | CRX037670 | CRA001315 | Wild type | MeRIP-seq | ThyroidGland |
| CRR042316 | CRX037672 | CRA001315 | Wild type | MeRIP-seq | Trachea |
| CRR042318 | CRX037674 | CRA001315 | Wild type | MeRIP-seq | UrinaryBladder |
| CRR042320 | CRX037676 | CRA001315 | Wild type | MeRIP-seq | UrinaryBladder |
| CRR055525 | CRX049951 | CRA001315 | Wild type | MeRIP-seq | Adipose |
| CRR055527 | CRX049953 | CRA001315 | Wild type | MeRIP-seq | Heart |
| CRR055529 | CRX049955 | CRA001315 | Wild type | MeRIP-seq | Spleen |
| CRR055533 | CRX049959 | CRA001315 | Wild type | MeRIP-seq | Lung |
| CRR055536 | CRX049961 | CRA001315 | Wild type | MeRIP-seq | Spleen |
| CRR055537 | CRX049963 | CRA001315 | Wild type | MeRIP-seq | Tongue |
| CRR055539 | CRX049965 | CRA001315 | Wild type | MeRIP-seq | UrinaryBladder |
| CRR055541 | CRX049967 | CRA001315 | Wild type | MeRIP-seq | Spleen |
| CRR055545 | CRX049971 | CRA001315 | Wild type | MeRIP-seq | Colon |
| CRR055547 | CRX049973 | CRA001315 | Wild type | MeRIP-seq | Esophagus |
| CRR055549 | CRX049975 | CRA001315 | Wild type | MeRIP-seq | Muscle |
| CRR055553 | CRX049979 | CRA001315 | Wild type | MeRIP-seq | Cerebrum |
| CRR055559 | CRX049985 | CRA001315 | Wild type | MeRIP-seq | Rectum |
| CRR055561 | CRX049987 | CRA001315 | Wild type | MeRIP-seq | Esophagus |
| CRR055563 | CRX049989 | CRA001315 | Wild type | MeRIP-seq | Rectum |
| CRR072991 | CRX062315 | CRA001315 | Wild type | MeRIP-seq | GOS |
| CRR072993 | CRX062317 | CRA001315 | Wild type | MeRIP-seq | GOS |
| CRR072999 | CRX062323 | CRA001315 | Wild type | MeRIP-seq | HT29 |
| CRR073005 | CRX062329 | CRA001315 | Wild type | MeRIP-seq | U251 |
| CRR073016 | CRX062340 | CRA001315 | Wild type | MeRIP-seq | Cerebellum |
| CRR073018 | CRX062342 | CRA001315 | Wild type | MeRIP-seq | Lung |
| CRR073020 | CRX062344 | CRA001315 | Wild type | MeRIP-seq | Lung |
| SRR10149738 | GSM4086538 | GSE137752 | Wild type | MeRIP-seq | ImmortalizedMyelogenousLeukemia |
| SRR10149740 | GSM4086540 | GSE137752 | Wild type | MeRIP-seq | ImmortalizedMyelogenousLeukemia |
| SRR10690133 | GSM4217473 | GSE141993 | Wild type | MeRIP-seq | IMR90 |
| SRR10690134 | GSM4217474 | GSE141993 | Wild type | MeRIP-seq | IMR90 |
| SRR11575353 | GSM4486700 | GSE148962 | Wild type | MeRIP-seq | HTR8SVneo |
| SRR11575354 | GSM4486701 | GSE148962 | Wild type | MeRIP-seq | HTR8SVneo |

| | | | | | |
|---|---|---|---|---|---|
| SRR11575355 | GSM4486702 | GSE148962 | Perturbation | MeRIP-seq | HTR8SVneo |
| SRR11575356 | GSM4486703 | GSE148962 | Perturbation | MeRIP-seq | HTR8SVneo |
| SRR11626645 | GSM4504165 | GSE149510 | Perturbation | MeRIP-seq | HCCLM3 |
| SRR11626647 | GSM4504167 | GSE149510 | Perturbation | MeRIP-seq | HCCLM3 |
| SRR11626649 | GSM4504169 | GSE149510 | Wild type | MeRIP-seq | HCCLM3 |
| SRR11626651 | GSM4504171 | GSE149510 | Wild type | MeRIP-seq | HCCLM3 |
| SRR11816543 | GSM4560324 | GSE150892 | Wild type | MeRIP-seq | HumanBLymphocyteP493_6 |
| SRR1182583 | GSM1339393 | GSE55572 | Wild type | MeRIP-seq | HumanNeuralProgenitor |
| SRR1182584 | GSM1339394 | GSE55572 | Wild type | MeRIP-seq | HumanNeuralProgenitor |
| SRR1182587 | GSM1339395 | GSE55572 | Wild type | MeRIP-seq | HumanNeuralProgenitor |
| SRR1182588 | GSM1339396 | GSE55572 | Wild type | MeRIP-seq | HumanNeuralProgenitor |
| SRR1182591 | GSM1339397 | GSE55572 | Perturbation | MeRIP-seq | HEK293T |
| SRR1182593 | GSM1339399 | GSE55572 | Perturbation | MeRIP-seq | HEK293T |
| SRR1182595 | GSM1339401 | GSE55572 | Wild type | MeRIP-seq | HEK293T |
| SRR1182603 | GSM1339409 | GSE55572 | Perturbation | MeRIP-seq | A549 |
| SRR1182605 | GSM1339411 | GSE55572 | Perturbation | MeRIP-seq | A549 |
| SRR1182607 | GSM1339413 | GSE55572 | Perturbation | MeRIP-seq | A549 |
| SRR1182609 | GSM1339415 | GSE55572 | Perturbation | MeRIP-seq | A549 |
| SRR1182611 | GSM1339417 | GSE55572 | Perturbation | MeRIP-seq | A549 |
| SRR1182613 | GSM1339419 | GSE55572 | Perturbation | MeRIP-seq | A549 |
| SRR1182615 | GSM1339421 | GSE55572 | Perturbation | MeRIP-seq | A549 |
| SRR1182617 | GSM1339423 | GSE55572 | Perturbation | MeRIP-seq | A549 |
| SRR1182619 | GSM1339425 | GSE55572 | Perturbation | MeRIP-seq | A549 |
| SRR1182621 | GSM1339427 | GSE55572 | Perturbation | MeRIP-seq | A549 |
| SRR1182623 | GSM1339429 | GSE55572 | Perturbation | MeRIP-seq | A549 |
| SRR1182627 | GSM1339433 | GSE55572 | Perturbation | MeRIP-seq | A549 |
| SRR1182631 | GSM1339437 | GSE55572 | Perturbation | MeRIP-seq | A549 |
| SRR1182633 | GSM1339439 | GSE55572 | Wild type | MeRIP-seq | A549 |
| SRR12235501 | GSM4673360 | GSE154555 | Wild type | MeRIP-seq | K180 |
| SRR12235502 | GSM4673361 | GSE154555 | Perturbation | MeRIP-seq | K180 |
| SRR12518106 | GSM4746128 | GSE156886 | Wild type | MeRIP-seq | Ameloblastoma |
| SRR12518108 | GSM4746130 | GSE156886 | Wild type | MeRIP-seq | Ameloblastoma |
| SRR12518110 | GSM4746132 | GSE156886 | Wild type | MeRIP-seq | Ameloblastoma |
| SRR12518112 | GSM4746134 | GSE156886 | Wild type | MeRIP-seq | NormalOralTissue |
| SRR12518114 | GSM4746136 | GSE156886 | Wild type | MeRIP-seq | NormalOralTissue |
| SRR12518116 | GSM4746138 | GSE156886 | Wild type | MeRIP-seq | NormalOralTissue |

| | | | | | |
|---|---|---|---|---|---|
| SRR12820682 | GSM4831167 | GSE137752 | Wild type | MeRIP-seq | MicroglialCellLineHMC3 |
| SRR13084754 | GSM4914534 | GSE161792 | Wild type | MeRIP-seq | MRC5 |
| SRR13084756 | GSM4914536 | GSE161792 | Wild type | MeRIP-seq | MRC5 |
| SRR13084758 | GSM4914538 | GSE161792 | Perturbation | MeRIP-seq | MRC5 |
| SRR13084760 | GSM4914540 | GSE161792 | Perturbation | MeRIP-seq | MRC5 |
| SRR13084762 | GSM4914542 | GSE161792 | Perturbation | MeRIP-seq | MRC5 |
| SRR13084764 | GSM4914544 | GSE161792 | Perturbation | MeRIP-seq | MRC5 |
| SRR13084766 | GSM4914546 | GSE161792 | Perturbation | MeRIP-seq | MRC5 |
| SRR13084768 | GSM4914548 | GSE161792 | Perturbation | MeRIP-seq | MRC5 |
| SRR13091694 | GSM4916321 | GSE161879 | Wild type | MeRIP-seq | NormalSalivaryGland |
| SRR13091696 | GSM4916323 | GSE161879 | Wild type | MeRIP-seq | InvasiveMalignantPleomorphicAdenomaTissue |
| SRR13091698 | GSM4916325 | GSE161879 | Wild type | MeRIP-seq | NormalSalivaryGland |
| SRR13091700 | GSM4916327 | GSE161879 | Wild type | MeRIP-seq | InvasiveMalignantPleomorphicAdenomaTissue |
| SRR13091702 | GSM4916329 | GSE161879 | Wild type | MeRIP-seq | NormalSalivaryGland |
| SRR13091704 | GSM4916331 | GSE161879 | Wild type | MeRIP-seq | InvasiveMalignantPleomorphicAdenomaTissue |
| SRR13091706 | GSM4916333 | GSE161879 | Wild type | MeRIP-seq | NormalSalivaryGland |
| SRR13091708 | GSM4916335 | GSE161879 | Wild type | MeRIP-seq | InvasiveMalignantPleomorphicAdenomaTissue |
| SRR13091710 | GSM4916337 | GSE161879 | Wild type | MeRIP-seq | NormalSalivaryGland |
| SRR13091712 | GSM4916339 | GSE161879 | Wild type | MeRIP-seq | InvasiveMalignantPleomorphicAdenomaTissue |
| SRR13948593 | GSM5169206 | GSE168778 | Wild type | MeRIP-seq | Kasumi_1 |
| SRR13948593 | GSM5169206 | GSE168778 | Perturbation | MeRIP-seq | Kasumi_1 |
| SRR13948594 | GSM5169207 | GSE168778 | Wild type | MeRIP-seq | Kasumi_1 |
| SRR13948594 | GSM5169207 | GSE168778 | Perturbation | MeRIP-seq | Kasumi_1 |
| SRR13948595 | GSM5169208 | GSE168778 | Wild type | MeRIP-seq | Kasumi_1 |
| SRR13948595 | GSM5169208 | GSE168778 | Perturbation | MeRIP-seq | Kasumi_1 |
| SRR13948596 | GSM5169209 | GSE168779 | Perturbation | MeRIP-seq | Kasumi_1 |
| SRR13948597 | GSM5169210 | GSE168780 | Perturbation | MeRIP-seq | Kasumi_1 |
| SRR13948598 | GSM5169211 | GSE168781 | Perturbation | MeRIP-seq | Kasumi_1 |

| | | | | | |
|---|---|---|---|---|---|
| SRR14134651 | GSM5224621 | GSE171373 | Wild type | MeRIP-seq | HUCMSCs |
| SRR14134653 | GSM5224623 | GSE171373 | Wild type | MeRIP-seq | HUCMSCs |
| SRR14134655 | GSM5224625 | GSE171373 | Perturbation | MeRIP-seq | HUCMSCs |
| SRR14134657 | GSM5224627 | GSE171373 | Wild type | MeRIP-seq | HUCMSCs |
| SRR14134659 | GSM5224629 | GSE171373 | Wild type | MeRIP-seq | HUCMSCs |
| SRR14134661 | GSM5224631 | GSE171373 | Wild type | MeRIP-seq | HUCMSCs |
| SRR14142650 | GSM5226075 | GSE171472 | Wild type | MeRIP-seq | H322 |
| SRR14142652 | GSM5226077 | GSE171472 | Perturbation | MeRIP-seq | H322 |
| SRR14765569 | GSM5365386 | GSE151028 | IVT | MeRIP-seq | HEK293T |
| SRR14765571 | GSM5365388 | GSE151028 | IVT | MeRIP-seq | HEK293T |
| SRR14765577 | GSM5365394 | GSE151028 | IVT | MeRIP-seq | HEK293T |
| SRR14765579 | GSM5365396 | GSE151028 | IVT | MeRIP-seq | HEK293T |
| SRR14765585 | GSM5365402 | GSE151028 | IVT | MeRIP-seq | HEK293T |
| SRR14765587 | GSM5365404 | GSE151028 | IVT | MeRIP-seq | HEK293T |
| SRR5080307 | GSM2417477 | GSE90914 | Wild type | MeRIP-seq | HEK293A |
| SRR5080308 | GSM2417478 | GSE90914 | Wild type | MeRIP-seq | HEK293A |
| SRR5080309 | GSM2417479 | GSE90914 | Wild type | MeRIP-seq | HEK293A |
| SRR5080310 | GSM2417480 | GSE90914 | Perturbation | MeRIP-seq | HEK293A |
| SRR5080311 | GSM2417481 | GSE90914 | Perturbation | MeRIP-seq | HEK293A |
| SRR5080312 | GSM2417482 | GSE90914 | Perturbation | MeRIP-seq | HEK293A |
| SRR6211496 | GSM2830065 | GSE106122 | Wild type | MeRIP-seq | HEL |
| SRR6211497 | GSM2830066 | GSE106122 | Wild type | MeRIP-seq | HEL |
| SRR7075097 | GSM3119587 | GSE113798 | Wild type | MeRIP-seq | HealthyMale |
| SRR7075098 | GSM3119588 | GSE113798 | Wild type | MeRIP-seq | HealthyMale |
| SRR7075099 | GSM3119589 | GSE113798 | Wild type | MeRIP-seq | MDDDiagnosed Male |
| SRR7075100 | GSM3119590 | GSE113798 | Wild type | MeRIP-seq | MDDDiagnosed Male |
| SRR7075101 | GSM3119591 | GSE113798 | Wild type | MeRIP-seq | HealthyMale |
| SRR7075102 | GSM3119592 | GSE113798 | Wild type | MeRIP-seq | HealthyMale |
| SRR7075103 | GSM3119593 | GSE113798 | Wild type | MeRIP-seq | MDDDiagnosed Male |
| SRR7075104 | GSM3119594 | GSE113798 | Wild type | MeRIP-seq | MDDDiagnosed Male |
| SRR7075105 | GSM3119595 | GSE113798 | Wild type | MeRIP-seq | HealthyMale |
| SRR7075106 | GSM3119596 | GSE113798 | Wild type | MeRIP-seq | HealthyMale |
| SRR7075107 | GSM3119597 | GSE113798 | Wild type | MeRIP-seq | MDDDiagnosed Male |
| SRR7075108 | GSM3119598 | GSE113798 | Wild type | MeRIP-seq | MDDDiagnosed Male |
| SRR7763571 | GSM3359937 | GSE119168 | Wild type | MeRIP-seq | OmentalTumor |
| SRR7763572 | GSM3359938 | GSE119168 | Wild type | MeRIP-seq | OmentalTumor |
| SRR7763573 | GSM3359939 | GSE119168 | Wild type | MeRIP-seq | OmentalTumor |
| SRR7763574 | GSM3359940 | GSE119168 | Wild type | MeRIP-seq | OmentalTumor |

| | | | | | |
|---|---|---|---|---|---|
| SRR7763575 | GSM3359941 | GSE119168 | Wild type | MeRIP-seq | OmentalTumor |
| SRR7763576 | GSM3359942 | GSE119168 | Wild type | MeRIP-seq | OmentalTumor |
| SRR7763577 | GSM3359943 | GSE119168 | Wild type | MeRIP-seq | NormalFallopian Tube |
| SRR7763578 | GSM3359944 | GSE119168 | Wild type | MeRIP-seq | NormalFallopian Tube |
| SRR7763579 | GSM3359945 | GSE119168 | Wild type | MeRIP-seq | NormalFallopian Tube |
| SRR7763580 | GSM3359946 | GSE119168 | Wild type | MeRIP-seq | NormalFallopian Tube |
| SRR7763581 | GSM3359947 | GSE119168 | Wild type | MeRIP-seq | NormalFallopian Tube |
| SRR7763582 | GSM3359948 | GSE119168 | Wild type | MeRIP-seq | NormalFallopian Tube |
| SRR7763583 | GSM3359949 | GSE119168 | Wild type | MeRIP-seq | NormalFallopian Tube |
| SRR7829546 | GSM3389537 | GSE119963 | Wild type | MeRIP-seq | ParentalPEO1 |
| SRR7829547 | GSM3389538 | GSE119963 | Perturbation | MeRIP-seq | ParentalPEO1 |
| SRR7881527 | GSM3396437 | GSE120229 | Wild type | MeRIP-seq | HEK293T |
| SRR7881528 | GSM3396438 | GSE120229 | Wild type | MeRIP-seq | HEK293T |
| SRR7881529 | GSM3396439 | GSE120229 | Perturbation | MeRIP-seq | HEK293T |
| SRR7881530 | GSM3396440 | GSE120229 | Perturbation | MeRIP-seq | HEK293T |
| SRR8209837 | GSM3484487 | GSE122744 | Wild type | MeRIP-seq | Cerebellumrip |
| SRR8209839 | GSM3484489 | GSE122744 | Wild type | MeRIP-seq | Cerebellumrip |
| SRR8209841 | GSM3484491 | GSE122744 | Wild type | MeRIP-seq | Cerebellumrip |
| SRR8209843 | GSM3484493 | GSE122744 | Wild type | MeRIP-seq | FrotalCortex |
| SRR8209845 | GSM3484495 | GSE122744 | Wild type | MeRIP-seq | FrotalCortex |
| SRR8209847 | GSM3484497 | GSE122744 | Wild type | MeRIP-seq | FrotalCortex |
| SRR8209849 | GSM3484499 | GSE122744 | Wild type | MeRIP-seq | Heart |
| SRR8209853 | GSM3484501 | GSE122744 | Wild type | MeRIP-seq | Heart |
| SRR8209855 | GSM3484503 | GSE122744 | Wild type | MeRIP-seq | Heart |
| SRR8209857 | GSM3484505 | GSE122744 | Wild type | MeRIP-seq | Kidney |
| SRR8209859 | GSM3484507 | GSE122744 | Wild type | MeRIP-seq | Kidney |
| SRR8209861 | GSM3484509 | GSE122744 | Wild type | MeRIP-seq | Kidney |
| SRR8209863 | GSM3484511 | GSE122744 | Wild type | MeRIP-seq | Liver |
| SRR8209865 | GSM3484513 | GSE122744 | Wild type | MeRIP-seq | Liver |
| SRR8209867 | GSM3484515 | GSE122744 | Wild type | MeRIP-seq | Liver |
| SRR8209869 | GSM3484517 | GSE122744 | Wild type | MeRIP-seq | Lung |
| SRR8209871 | GSM3484519 | GSE122744 | Wild type | MeRIP-seq | Lung |
| SRR8209873 | GSM3484521 | GSE122744 | Wild type | MeRIP-seq | Lung |
| SRR8209875 | GSM3484523 | GSE122744 | Wild type | MeRIP-seq | Muscle |
| SRR8209877 | GSM3484525 | GSE122744 | Wild type | MeRIP-seq | Muscle |
| SRR8209879 | GSM3484527 | GSE122744 | Wild type | MeRIP-seq | Muscle |
| SRR8209881 | GSM3484529 | GSE122744 | Wild type | MeRIP-seq | Spleen |

| | | | | | |
|---|---|---|---|---|---|
| SRR8209883 | GSM3484531 | GSE122744 | Wild type | MeRIP-seq | Spleen |
| SRR8209885 | GSM3484533 | GSE122744 | Wild type | MeRIP-seq | Spleen |
| SRR8742553 | GSM3675280 | GSE128443 | Wild type | MeRIP-seq | HeLa |
| SRR8742555 | GSM3675282 | GSE128443 | Perturbation | MeRIP-seq | HeLa |
| SRR8742557 | GSM3675284 | GSE128443 | Wild type | MeRIP-seq | HeLa |
| SRR8742559 | GSM3675286 | GSE128443 | Perturbation | MeRIP-seq | HeLa |
| SRR8742561 | GSM3675288 | GSE128443 | Wild type | MeRIP-seq | HeLa |
| SRR8742563 | GSM3675290 | GSE128443 | Perturbation | MeRIP-seq | HeLa |
| SRR8755805 | GSM3678928 | GSE128575 | Wild type | MeRIP-seq | THP1 |
| SRR8755806 | GSM3678929 | GSE128575 | Wild type | MeRIP-seq | THP1 |
| SRR8755807 | GSM3678930 | GSE128575 | Perturbation | MeRIP-seq | THP1 |
| SRR8755808 | GSM3678931 | GSE128575 | Perturbation | MeRIP-seq | THP1 |
| SRR8755809 | GSM3678932 | GSE128575 | Perturbation | MeRIP-seq | THP1 |
| SRR8755810 | GSM3678933 | GSE128575 | Perturbation | MeRIP-seq | THP1 |
| SRR8889196 | GSM3720795 | GSE129716 | Wild type | MeRIP-seq | HCT116 |
| SRR8889198 | GSM3720797 | GSE129716 | Wild type | MeRIP-seq | HCT116 |
| SRR8889201 | GSM3720780 | GSE129716 | Perturbation | MeRIP-seq | HCT116 |
| SRR8889203 | GSM3720782 | GSE129716 | Perturbation | MeRIP-seq | HCT116 |
| SRR9336433 | GSM3900733 | GSE133132 | Perturbation | MeRIP-seq | BGC823 |
| SRR9336434 | GSM3900734 | GSE133132 | Wild type | MeRIP-seq | BGC823 |
| SRR9974311 | GSM4037123 | GSE106122 | Wild type | MeRIP-seq | HEL |
| SRR9974312 | GSM4037124 | GSE106122 | Wild type | MeRIP-seq | HEL |
| SRR9974313 | GSM4037125 | GSE106122 | Wild type | MeRIP-seq | HEL |
| SRR9974315 | GSM4037127 | GSE106122 | Wild type | MeRIP-seq | ERY2 |
| SRR9974317 | GSM4037129 | GSE106122 | Wild type | MeRIP-seq | CD14 |
| SRR9974319 | GSM4037131 | GSE106122 | Wild type | MeRIP-seq | MEG |
| SRR9974321 | GSM4037133 | GSE106122 | Wild type | MeRIP-seq | ERY1 |
| SRR9974323 | GSM4037135 | GSE106122 | Wild type | MeRIP-seq | CMP |
| SRR9974325 | GSM4037137 | GSE106122 | Wild type | MeRIP-seq | ERY3 |
| SRR9974327 | GSM4037139 | GSE106122 | Wild type | MeRIP-seq | GMP |
| SRR9974329 | GSM4037141 | GSE106122 | Wild type | MeRIP-seq | MEP |
| SRR9974331 | GSM4037143 | GSE106122 | Wild type | MeRIP-seq | HSC |
| ERR11278802 | ERS6364536 | PRJEB44348 | Wild type | Oxford-nanopore | A549 |
| ERR5867843 | ERS6364498 | PRJEB44348 | Wild type | Oxford-nanopore | HCT116 |
| ERR5867844 | ERS6364544 | PRJEB44348 | Wild type | Oxford-nanopore | HCT116 |
| ERR5867845 | ERS6364499 | PRJEB44348 | Wild type | Oxford-nanopore | HCT116 |
| ERR5867847 | ERS6364500 | PRJEB44348 | Wild type | Oxford-nanopore | HCT116 |
| ERR9286473 | ERS6364541 | PRJEB44348 | Wild type | Oxford-nanopore | A549 |
| ERR9286474 | ERS6364540 | PRJEB44348 | Wild type | Oxford-nanopore | A549 |
| ERR9286476 | ERS6364539 | PRJEB44348 | Wild type | Oxford-nanopore | A549 |
| ERR9286486 | ERS6364545 | PRJEB44348 | Wild type | Oxford-nanopore | HCT116 |
| ERR9286487 | ERS6364546 | PRJEB44348 | Wild type | Oxford-nanopore | HCT116 |
| ERR9286488 | ERS6364547 | PRJEB44348 | Wild type | Oxford-nanopore | HCT116 |
| ERR9286489 | ERS6364548 | PRJEB44348 | Wild type | Oxford-nanopore | HCT116 |

| | | | | | |
|---|---|---|---|---|---|
| ERR9286490 | ERS6364549 | PRJEB44348 | Wild type | Oxford-nanopore | HCT116 |
| ERR9286491 | ERS6364550 | PRJEB44348 | Wild type | Oxford-nanopore | HCT116 |
| ERR9286492 | ERS6364556 | PRJEB44348 | Wild type | Oxford-nanopore | HCT116 |
| ERR9286493 | ERS6364557 | PRJEB44348 | Wild type | Oxford-nanopore | HCT116 |
| ERR9286495 | ERS6597012 | PRJEB44348 | Wild type | Oxford-nanopore | HEK293T |
| ERR9286496 | ERS6597013 | PRJEB44348 | Wild type | Oxford-nanopore | HEK293T |
| ERR9286497 | ERS6597014 | PRJEB44348 | Wild type | Oxford-nanopore | HEK293T |
| ERR9286498 | ERS6597015 | PRJEB44348 | Wild type | Oxford-nanopore | HEK293T |

### Supplementary Methodological Details

**Content:**


I. Introduction
II. Data Processing
    **1. Data gathering**
    2. Quality Control
    3. Alignment
    4. Feature counting
    5. Omics data frameworks
    5. IVT calibration
III. Site-calling and Normalization: Beta-binomial mixture model
    **Beta-binomial mixture model**
    **Expectation-Maximization (EM) algorithm**
        1. Initialization
        2. E-step
        3. M-step
        4. Convergence
    Model outputs
IV. Technical Orthogonal Integration: IDR analysis
    IDR analysis
        **1. Site ranking**
        **2. Modelling**
        **3. Threshold selection**
    Technical orthogonal integration
V. Codes
VI. References

### I. Introduction

The m6AConquer database offers curated quantitative m6A profiling sequencing data sets. This page provides a brief overview of the processing workflows utilized to convert raw reads into the data available in this database.

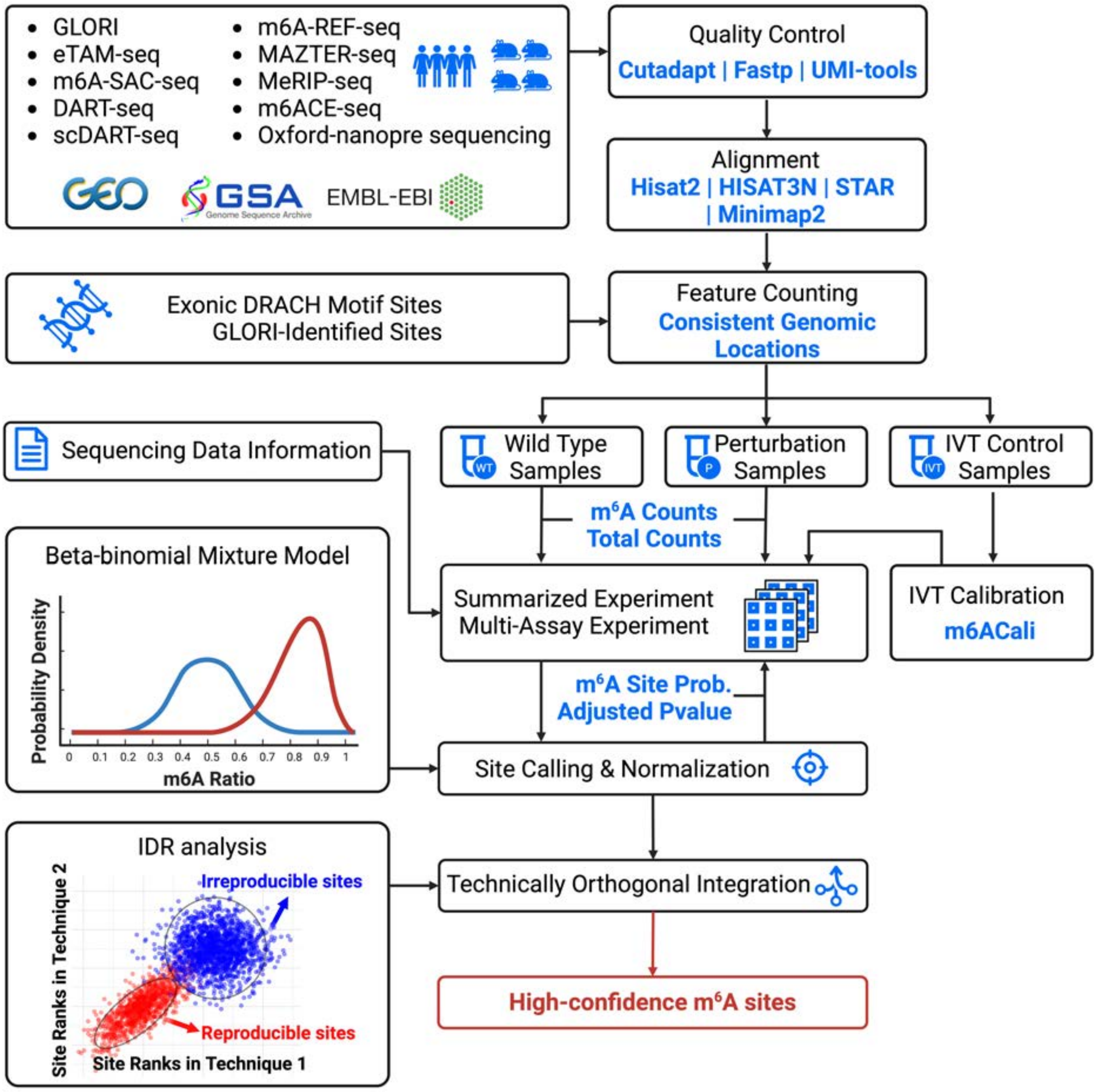


The workflow of m6AConquer

### II. Data Processing

#### 1. Data gathering

m6AConquer summarises sequencing datasets from 10 different m6A profiling techniques. Both wild type and perturbation samples are included in the database. The perturbation types mainly include the manipulation of m6A-related genes, such as the over-expression of FTO and the knockout of Mettl3. IVT control samples are also included for quantification calibration.

All the sequencing datasets are downloaded from three central database (GEO, GSA, and EMBL-EBI).

#### 2. Quality Control

FastQ read files for NGS techniques are retrieved and downloaded accordingly. Most reads undergo initial trimming using tools like Cutadapt, Trim Galore, and Flexbar to eliminate adaptors and low-quality reads. Some reads are deduplicated through Samtools, Fastx Collapser, and UMI-tools to remove PCR duplications. Fastp is used for both trimming and deduplication.

As for Oxford Nanopore sequencing data, the raw Fast5 files are obtained from the Singapore Nanopore-Expression Project. Quality control is conducted during the basecalling process with Guppy.

The table below summarize the software and tools used in data cleaning and read mapping:

| Techniques | Data cleaning | Read mapping |
|---|---|---|
| DART-seq | Fastp v.0.23.4 | HISAT-3N v.2.2.1-3n-0.0.3 (C → T) |
| eTAM-seq | Cutadapt v.2.9, UMI-tools v.1.1.4 | HISAT-3N v.2.2.1-3n-0.0.3 (A → G) |
| GLORI | Trim Galore v.0.6.6, Fastp v.0.23.4 | STAR v.2.7.11a, bowtie v.1.3.1 |
| m6A-REF-seq | Cutadapt v.2.9 | HISAT2 v.2.1.0 |
| m6A-SAC-seq | Cutadapt v.2.9, Fastx_collapser v.0.0.13 | STAR v.2.7.11a |
| m6ACE | Fastp v.0.23.4 | STAR v.2.7.11a |
| MAZTER-seq | Trim Galore v.0.6.6 | STAR v.2.7.11a |
| MeRIP-seq | Trim Galore v.0.6.6 | HISAT2 v.2.1.0 |

| Techniques | Data cleaning | Read mapping |
|---|---|---|
| Oxford-nanopore | Guppy v.3.1.5 (Basecalling) | Minimap2 v.2.17 |
| scDART-seq | Flexbar | STAR v.2.7.11a |

#### 3. Alignment

For NGS sequencing data, 4 aligners (STAR, Bowtie, HISAT2, and HISAT-3N) are utilized for different techniques based on their original pipelines. Bowtie is employed explicitly for transcriptome alignment, while the other three aligners are utilized for genome alignment. The genome index of HISAT2 can be obtained from the provided link: https://genome-idx.s3.amazonaws.com/hisat/grch38_genome.tar.gz. Meanwhile, the genome indexes for STAR and HISAT-3N are constructed as per the specific requirements. All sequencing reads are aligned to the GRCh38 or hg38 human genome, and various versions of gene annotation from RefSeq, Genecode, and Ensemble are employed under their original pipelines. The codes for index building can be accessed on GitHub.

In the context of Oxford Nanopore sequencing data, the aligner minimap2 is utilized for transcriptome alignment. The Ensemble transcript reference of GRCh38 is indexed using the default parameters of minimap2.

#### 4. Feature counting

The m6AConquer database summarizes quantitative m6A profiling results in the form of SummarizedExperiment (SE) objects. These SE objects summarize the total and m6A read coverages at a consistent range of genomic locations and gene expression levels. The consistent genomic locations consist of 2526220 adenosines (A) at the center of exonic DRACH motifs and 262485 GLORI m6A sites discovered at other genomic locations reported in the supplementary data of GLORI. The m6AConquer database also includes gene expression matrices in SE objects on 58560 genes (Transctipt-Omics).

#### 5. Omics data frameworks

The SummarizedExperiment objects consist of four primary components: row ranges, column data, assays matrices, and metadata.

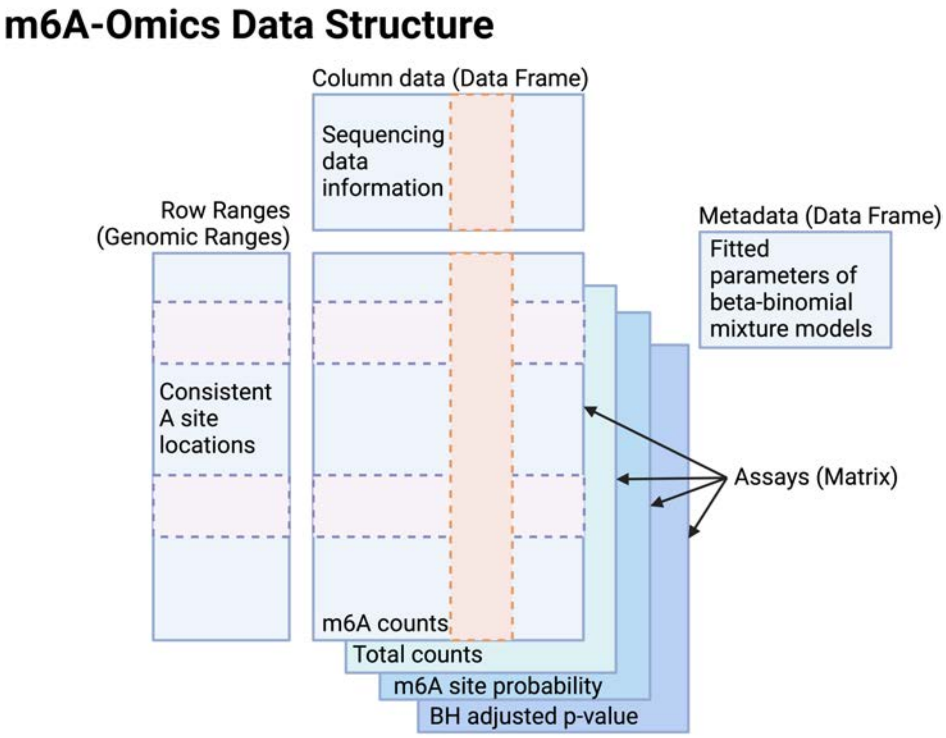


Data structure of SummarizedExperiment objects for m6A-Omics.

In SE objects for m6A-omics,

- **Assays**: The matrices for m6A counts and total read coverages at this candidate A sites, m6A site probabilities predicted by beta-binomial mixture models, and the Benjamini–Hochberg method corrected p-values (FDR).
- **Row ranges**: Genomic ranges of consistent A locations.
- **Column data**: Sequencing data information provided by the data generators in the public repositories.
- **Metadata**: Fitted parameters of beta-binomial mixture models and IDR outcomes.

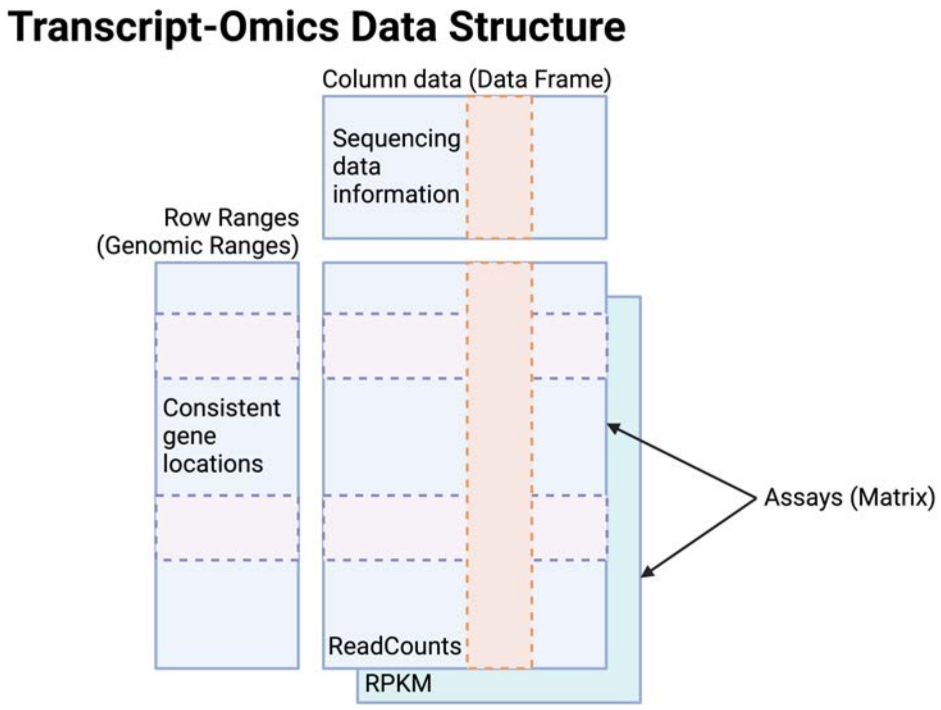


Data structure of SummarizedExperiment objects for Transcript-Omics.

In SE objects for transcript-omics,

- **Assays**: The matrices for read counts and RPKM at consistent gene features.
- **Row ranges**: Genomic ranges of gene features.
- **Column data**: Sequencing data information provided by the data generators in the public repositories, identical to m6A-Omics SE.

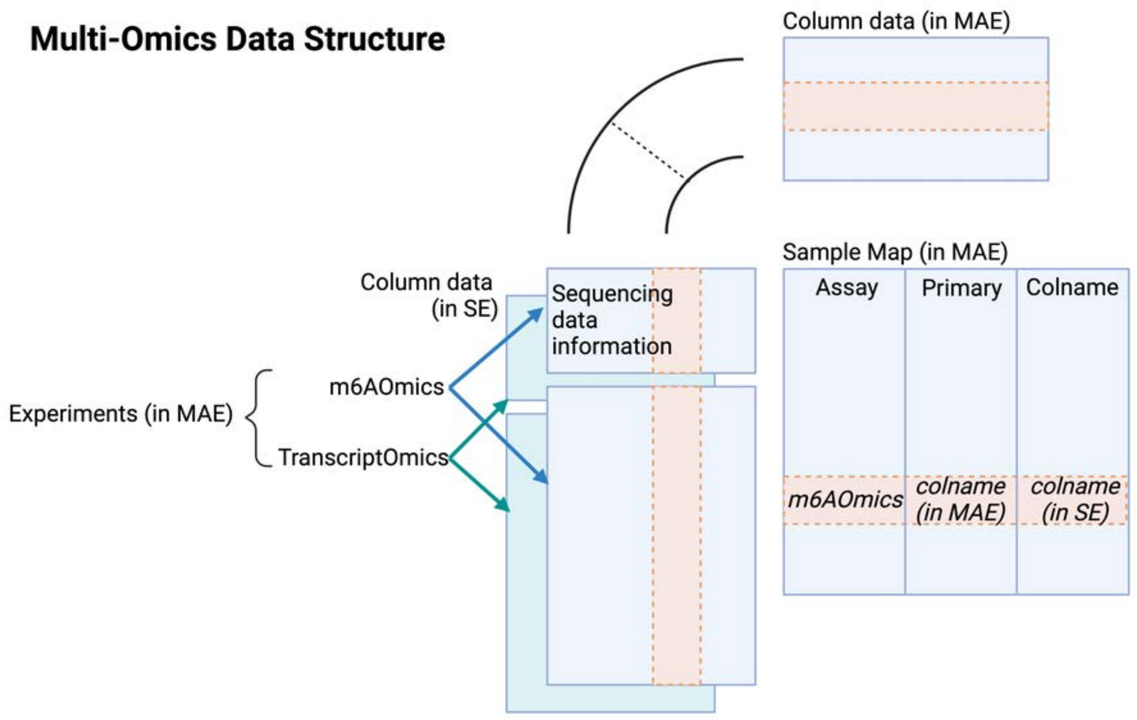


Data structure of Multi-Assay Experiment objects for Multi-Omics.

Apart from these single-omics data in SE format, multi-omics data in the form of MultiAssayExperiment (MAE) for each profiling technique are also provided. The MAE files are the integration of m6A-Omics and Transcrit-Omics SE objects. The MAE contains distinct components such as experiments, sample maps, and column data for MAE.

In MAE objects for multi-omits,

- **Experiments:** m6A-Omics and Transcrit-Omics SE objects
- **Column data in MAE:** The union of column data in all experiment SEs.
- **Sample Map:** The map relating the column data in experiments to the column data in MAE.

All three types of files (multi-omics, m6A-omics, and transcript-omics) can all be downloaded from the Download page.

#### 5. IVT calibration

In the m6AConquer, we have incorporated in vitro transcription (IVT) control samples from GLORI, eTAM-seq, and MeRIP-seq datasets. The IVT control sequencing data from GLORI and eTAM-seq employ a beta-binomial mixture model to call m6A sites separately. Sites with a false discovery rate (FDR) below 0.05 are considered false positive m6A sites for the respective GLORI or eTAM-seq datasets. In the case of IVT control sequencing data from MeRIP-seq, m6ACali is used to identify false positive m6A sites. For other sequencing data within these three datasets, the m6A counts and total counts at their identified false positive m6A sites are set to 0.

### III. Site-calling and Normalization: Beta-binomial mixture model

#### Beta-binomial mixture model

The two-component beta-binomial mixture model is a statistical model that assumes the data is generated from a mixture of two beta distributions. Each beta distribution represents a different underlying mechanism or subgroup within the population. The probability of observing success for each individual is assumed to come from one of the two beta distributions, and the choice of distribution is determined by a latent (unobserved) variable that follows a binomial distribution:

$$x \sim \pi \cdot \mathrm{Beta}(\alpha_1, \beta_1) + (1 - \pi) \cdot \mathrm{Beta}(\alpha_2, \beta_2) \tag{1}$$

where $\pi$ and $1 - \pi$ are the proportion of two beta distribution components, and $\alpha$ and $\beta$ are beta distribution parameters.

#### Expectation-Maximization (EM) algorithm

In m6AConquer, we model the m6A methylation ratios detected in each sequencing sample as a two-component beta-binomial mixture distribution representing two possible methylation states: methylated state (foreground, "$\mathrm{fg}$") and unmethylated (background, "$\mathrm{bg}$") state. The fitting of the beta-binomial mixture model follows the Expectation-Maximization (EM) algorithm. The EM algorithm is an iterative method that starts with initial estimates for the parameters and then changes between the E-step (Expectation) and the M-step (Maximization). In the E-step, the log-likelihood is computed the  based on the current parameter estimates. In the M-step, the log-likelihood is maximized to update the parameter estimates.

#### 1. Initialization

For each site with m6A read coverages, let $m_i$ and $n_i$ be the m6A and total read coverages at site $i$. Let $\{x_i = \frac{m_i}{n_i}\},\ i = 1, 2 \ldots, N$, denotes the m6A methylation ratios at site $i$, where $N$ is the number of sites with m6A read coverage.

To obtain a closer initial estimate for $\pi$, a two-component binomial mixture distribution is fitted for $x_i$:

$$x \sim \pi \cdot \mathrm{Binomial}(N, p_{\mathrm{fg}}) + (1 - \pi) \cdot \mathrm{Binomial}(N, p_{\mathrm{bg}}) \tag{2}$$

The fitting of this primitive binomial mixture distribution is also implemented by EM algorithm.

To obtain initial estimates for $\alpha_{\mathrm{fg}}$, $\beta_{\mathrm{fg}}$, $\alpha_{\mathrm{bg}}$ , and $\beta_{\mathrm{bg}}$, weighted method of moments is firstly used for estimation (18-21), following the weighted maximum likelihood estimation (8-17) to obtain a closer estimation. The details of these two estimation methods are introduced in the M-step.

#### 2. E-step

In the E-step, the algorithm calculates the value of the likelihood function concerning the conditional distribution of the latent variable given the observed data and current estimate of the parameters. This is where posterior probabilities are computed:

$$\begin{aligned} r_{i,\mathrm{fg}} &= \frac{p(m_i|n_i, \alpha_{\mathrm{fg}}, \beta_{\mathrm{fg}}) \cdot p(\alpha_{\mathrm{fg}}, \beta_{\mathrm{fg}})}{p(m_i|\theta)} \\ &= \frac{p(m_i|n_i, \alpha_{\mathrm{fg}}, \beta_{\mathrm{fg}}) \cdot \pi}{p(m_i|n_i, \alpha_{\mathrm{fg}}, \beta_{\mathrm{fg}}) \cdot \pi + p(m_i|n_i, \alpha_{\mathrm{bg}}, \beta_{\mathrm{bg}}) \cdot (1 - \pi)} \end{aligned} \tag{3}$$

$$\underset{j \in \{\mathrm{fg}, \mathrm{bg}\}}{p(m_i|n_i, \alpha_j, \beta_j)} = \binom{n_i}{m_i} \frac{\mathrm{B}(m_i + \alpha_j, n_i - m_i + \beta_j)}{\mathrm{B}(\alpha_j, \beta_j)} \tag{4}$$

$$r_{i,\mathrm{bg}} = 1 - r_{i,\mathrm{fg}} \tag{5}$$

where $r_{i,\mathrm{fg}}$ and $r_{i,\mathrm{bg}}$ denotes the posterior probability of $x_i$ belonging to the foreground and background component respectively, $m_i$ and $n_i$ be the m6A and total read coverages, (4) denotes the probability mass function of beta distribution component, and $\mathrm{B}$ denotes beta function.

#### 3. M-step

In the M-step, the algorithm finds the parameters that maximize the likelihood. By default, weighted maximum likelihood estimation (WMLE) is used to obtained the estimated parameters. If the WMLE fails to converge, weighted method of moments is used in current iteration as the estimation method.

- **Update component proportion**

  The update formula for component proportions are:

$$\pi = \frac{\sum_{i=1}^{N} r_{i,\text{fg}}}{N} \tag{6}$$

$$\pi_j = \begin{cases} \pi & j = \text{fg} \\ 1 - \pi & j = \text{bg} \end{cases} \tag{7}$$

  where $N$ is the number of sites with m6A read coverage.

- **Update beta distribution parameters**

  - **Weighted maximum likelihood estimation (WMLE)**

    To avoid underflow, the log-likelihood of beta-binomial mixture distribution is calculated:

$$l = \sum_{i=1}^{n} \sum_{j \in \{\text{fg},\text{bg}\}} [\log \pi_j + \log p(m_i | n_i, \alpha_j, \beta_j)] \tag{8}$$

    To find the estimates of $\alpha$ and $\beta$ that maximize the log-likelihood function (8), the Newton-Raphson method is used. The Newton-Raphson method is an iterative root-finding algorithm that uses the first and second derivatives of a function to find the values of the parameters that maximize the function. The update formula in the Newton-Raphson method are:

$$parm_{\text{new}} = parm_{\text{old}} - Hessian_{\text{inv}} * f = \begin{bmatrix} \alpha_{\text{new}} \\ \beta_{\text{new}} \end{bmatrix} \tag{9}$$

$$Hessian = \begin{bmatrix} \frac{\partial^2 l}{\partial \alpha^2} & \frac{\partial^2 l}{\partial \alpha \partial \beta} \\ \\ \frac{\partial^2 l}{\partial \beta \partial \alpha} & \frac{\partial^2 l}{\partial \beta^2} \end{bmatrix} \tag{10}$$

$$f = \begin{bmatrix} \frac{\partial l}{\partial \alpha} \\ \\ \frac{\partial l}{\partial \beta} \end{bmatrix} \tag{11}$$

$$parm_{\text{old}} = \begin{bmatrix} \alpha_{\text{old}} \\ \\ \beta_{\text{old}} \end{bmatrix} \tag{12}$$

where $Hessian_{\text{inv}}$ is the inverse of Hessian matrix (10), $\frac{\partial l}{\partial \alpha}$ and $\frac{\partial l}{\partial \beta}$ are the first partial derivatives of (6) with respect to $\alpha_{fg}$ and $\beta_{fg}$ or $\alpha_{bg}$ and $\beta_{bg}$, and $\frac{\partial^2 l}{\partial \alpha^2}$, $\frac{\partial^2 l}{\partial \alpha \partial \beta}$ and $\frac{\partial^2 l}{\partial \beta^2}$ are the second partial derivatives. $\alpha_{\text{old}}$ and $\beta_{\text{old}}$ are $\alpha_{\text{fg}}$ and $\beta_{\text{fg}}$ or $\alpha_{\text{bg}}$ and $\beta_{\text{bg}}$ of the previous iteration, and $\alpha_{\text{new}}$ and $\beta_{\text{new}}$ are the updated parameters in current iteration.

The derivatives are calculates as follow:

$$\begin{aligned} \frac{\partial l}{\partial \alpha} = \sum_{i=1}^{N} & r_i(\psi(\alpha+\beta) - \psi(\alpha) \\ & + \psi(\alpha + m_i) - \psi(\alpha+\beta+n_i)) \end{aligned} \tag{13}$$

$$\begin{aligned} \frac{\partial l}{\partial \beta} = \sum_{i=1}^{N} & r_i(\psi(\alpha+\beta) - \psi(b) \\ & + \psi(\beta + m_i - x_i) - \psi(\alpha+\beta+n_i)) \end{aligned} \tag{14}$$

$$\begin{aligned} \frac{\partial^2 l}{\partial \alpha^2} = \sum_{i=1}^{N} & r_i(\psi_1(\alpha+\beta) - \psi_1(\alpha) \\ & + \psi_1(\alpha + m_i) - \psi_1(\alpha+\beta+n_i)) \end{aligned} \tag{15}$$

$$\begin{aligned} \frac{\partial^2 l}{\partial \beta^2} = \sum_{i=1}^{N} & r_i(\psi_1(\alpha+\beta) - \psi_1(\beta) \\ & + \psi_1(\beta + n_i - m_i) - \psi_1(\alpha+\beta+n_i)) \end{aligned} \tag{16}$$

$$\frac{\partial^2 l}{\partial\alpha\partial\beta} = \sum_{i=1}^{N} r_i(\psi_1(\alpha+\beta) - \psi_1(\alpha+\beta+n_i)) \tag{17}$$

where $\psi$ represents the digamma function, $\psi_1$ represents the trigamma function, $r_i$ are the responsibilities calculated in the E-step, $m_i$ are the m6A read coverage at site $i$, and $n_i$ are the total read coverage at site $i$.

The parameters for foreground and background components are estimated separately.

- **Weighted method of moments (WMM)**

  If WMLE fails to converge in a certain iteration, the estimated $\alpha$ and $\beta$ is substituted by those estimated by weighted method of moments (WMM):

$$\alpha_{\text{mom}} = \hat{\mu} \cdot \left( \frac{\hat{\mu} \cdot (1-\hat{\mu})}{\hat{\sigma}^2} - 1 \right) \tag{18}$$

$$\beta_{\text{mom}} = (1-\hat{\mu}) \cdot \left( \frac{\hat{\mu} \cdot (1-\hat{\mu})}{\hat{\sigma}^2} - 1 \right) \tag{19}$$

  where $\hat{\mu}$ is the weighted mean and $\hat{\sigma}^2$ is weighted variance:

$$\hat{\mu} = \frac{\sum_{i=1}^{N} r_i \cdot x_i}{\sum_{i=1}^{N} r_i} \tag{20}$$

$$\hat{\sigma}^2 = \frac{\sum_{i=1}^{N} r_i \cdot (x_i - \mu)^2}{\sum_{i=1}^{N} r_i} \tag{21}$$

  Here, the weight $r_i$ are the responsibilities calculated in the E-step.

  The parameters for foreground and background components are estimated separately.

#### 4. Convergence

The algorithm checks for convergence by testing if the absolute differences between the current and the previous parameter values are smaller than a certain tolerance. If not, the parameters are updated, and the next iteration starts. If yes, the process stops, and the final estimates for the parameters are returned.

### Model outputs

The beta-binomial mixture model in m6AConquer outputs two statistics: m6A site probabilities and false discovery rates.

- **m6A site probabilities**: These are the posterior probability (responsibilities) of the sites being assigned to the foreground component (3, 4) with the final fitted model. This can be regarded as the normalized m6A ratios.
- **False discovery rates (FDR)**: These are the Benjamini-Hochberg (BH) method corrected p-values. P-values are the probabilities of the true m6A counts equal or larger than the observed m6A counts in the background beta-binomial component. This can be calculated through the cumulative density function (CDF) of the beta-binomial distribution with the final fitted parameters for the background component:

$$\text{P-value} = 1 - F(x; \alpha_{bg}, \beta_{bg}) \tag{22}$$

$$F(x; \alpha_{bg}, \beta_{bg}) = f(0) + \cdots + f(x) \tag{23}$$

$$\begin{aligned} f(x) = & \prod_{k=1}^{x} (\frac{N+1-k}{k}) \\ & \cdot \frac{(\alpha_{bg} + x - 1)! \cdot (\beta_{bg} + N - x - 1)! \cdot \mathrm{B}(\alpha_{bg}, \beta_{bg})}{(\alpha_{bg} + \beta_{bg} + N + 1)!} \end{aligned} \tag{24}$$

where $x$ is the m6A counts, $F(x; \alpha_{bg}, \beta_{bg})$ and $f(x)$ is the cumulative density function (CDF) and probability mass function (PMF) of the background beta-binomial component respectively, $N$ is the number of sites with m6A read coverage and $\mathrm{B}$ is the beta function. The CDF of the background component is calculated through the sum of PMF, which can be implemented with a recursive algorithm.

### IV. Technical Orthogonal Integration: IDR analysis

#### IDR analysis

To identify reliable modification sites and investigate differences between techniques, m6AConquer implements Irreproducible Discovery Rate (IDR) analysis, a method commonly used in CHIP-seq data to identify consistent peaks in replicates. In our implementation, we evaluate the consistency of identified sites between different techniques, particularly those employing orthogonal mechanisms to quantify m6A abundances. IDR analysis involves three key steps to assess the reproducibility of methylation sites in various m6A profiling technique.

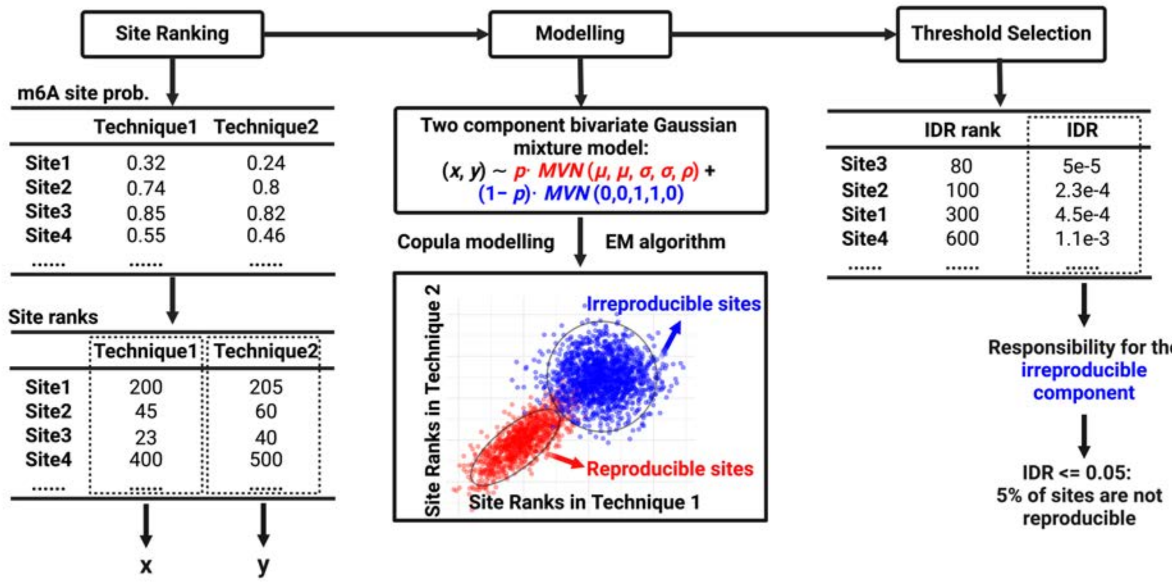


The workflow of IDR analysis

##### 1. Site ranking

The overlapped sites between each pair of profiling techniques are ranked based on the value of interest. In m6AConquer, m6A site probabilities are selected as the evaluation value. Therefore, the overlapped sites are ranked based on posterior probabilities, and the ranks are used as modelling statistics in the following step.

#### 2. Modelling

The underlying assumption in IDR analysis is that reproducibility sites should rank higher and consistently in each pair of techniques. To model the heterogeneity, a mixture framework is employed, where a positively correlated bivariate Gaussian distribution denotes the component of reproducible sites and a zero-correlated bivariate Gaussian distribution denotes the component of irreproducible sites.

$$\begin{aligned}(x, y) \sim & p \cdot \text{Normal}(\begin{pmatrix}\mu \\ \mu\end{pmatrix}, \begin{pmatrix}\sigma^2 & \rho \cdot \sigma^2 \\ \rho \cdot \sigma^2 & \sigma^2\end{pmatrix}) \\ & + (1-p) \cdot \text{Normal}(\begin{pmatrix}0 \\ 0\end{pmatrix}, \begin{pmatrix}1 & 0 \\ 0 & 1\end{pmatrix})\end{aligned} \tag{25}$$

where $(x, y)$ denotes site ranks of each pair of techniques, $p$ denotes the proportion of the reproducible site component, and $\mu$, $\sigma^2$, and $\rho$ denote the parameters for the positively correlated bivariate Gaussian distribution.

A two-component semiparametric copula mixture model is employed to assess the dependence structure between the ranks of paired techniques. The associations among the variables are modelled by a parametric joint distribution, while the marginal distributions are estimated nonparametrically based on their ranks. The Gaussian copula family is employed to model dependence structures, treating the observed data as being generated from underlying latent variables $(z_x, z_y)$. The underlying latent variables reflect the biological replicates and are assumed to have the same marginal distribution. Therefore, the joint parametric model for copula with latent variables $(z_x, z_y)$ is:

$$\begin{pmatrix}z_x \\ z_y\end{pmatrix} | K = k \sim \text{Normal}\left(\begin{pmatrix}\mu_k \\ \mu_k\end{pmatrix}, \begin{pmatrix}\sigma_k^2 & \rho_k \cdot \sigma_k^2 \\ \rho_k \cdot \sigma^2 & \sigma_k^2\end{pmatrix}\right), k = 0, 1 \tag{26}$$

where $K_i \sim \text{Bernoulli}(p)$, $\mu_0 = 0$, $\mu_1 > 0$, $\rho_0 = 0$, and $0 < \rho_1 < 1$.

The IDR value is the posterior probability that a rank pair $i$ denoted as $(x_i, y_i)$ belongs to the irreproducible component:

$$\text{IDR} = \frac{C_{\text{irrep}}}{C_{\text{irrep}} + C_{\text{rep}}} \tag{27}$$

$$C_{\text{irrep}} = p \cdot h_0(G^{-1}(F_x(x)), G^{-1}(F_y(y))) \tag{28}$$

$$C_{\text{rep}} = (1-p) \cdot h_1(G^{-1}(F_x(x)), G^{-1}(F_y(y))) \tag{29}$$

where $h_0 \sim \text{Normal}\left(\begin{pmatrix}0\\0\end{pmatrix}, \begin{pmatrix}1 & 0\\0 & 1\end{pmatrix}\right)$ and $h_1 \sim \text{Normal}\left(\begin{pmatrix}\mu_1\\\mu_1\end{pmatrix}, \begin{pmatrix}\sigma_1^2 & \rho_1 \cdot \sigma_1^2\\\rho_1 \cdot \sigma_1^2 & \sigma_1^2\end{pmatrix}\right)$. $F_x$ and $F_y$ are the unknown marginal distributions of ranks $x$ and $y$ respectively. $G^{-1}$ is are the right-continuous inverses of $G$ which is defined as:

$$u_x \equiv G(z_x) = \frac{p}{\sigma_1} \cdot \Phi\left(\frac{z_x - \mu_1}{\sigma_1}\right) + (1-p) \cdot \Phi(z_x) \tag{30}$$

$$u_y \equiv G(z_y) = \frac{p}{\sigma_1} \cdot \Phi\left(\frac{z_y - \mu_1}{\sigma_1}\right) + (1-p) \cdot \Phi(z_y) \tag{31}$$

$$x = F_x^{-1}(u_x) \tag{32}$$

$$y = F_y^{-1}(u_y) \tag{33}$$

where $F_x^{-1}$ and $F_y^{-1}$ are the right-continuous inverses of $F_x$ and $F_y$.

To fit procedure for this model is implemented with a EM algorithm coupled with "pseudo-likelihood" likelihood approach. For detailed fitting procedure, please refer to Li et al. (2011).

##### 3. Threshold selection

By setting a threshold for the IDR value, the irreproducible discoveries can be controlled. In m6AConquer, this threshold is set to 0.05, which means the overlapped sites with posterior probabilities of belonging to the irreproducible component of less than 5% are considered reproducible.

#### Technical orthogonal integration

IDR analysis is conducted between orthogonal technique pairs. Here, 10 m6A quantitative profiling techniques are classified into 4 groups based on their sequencing principles:

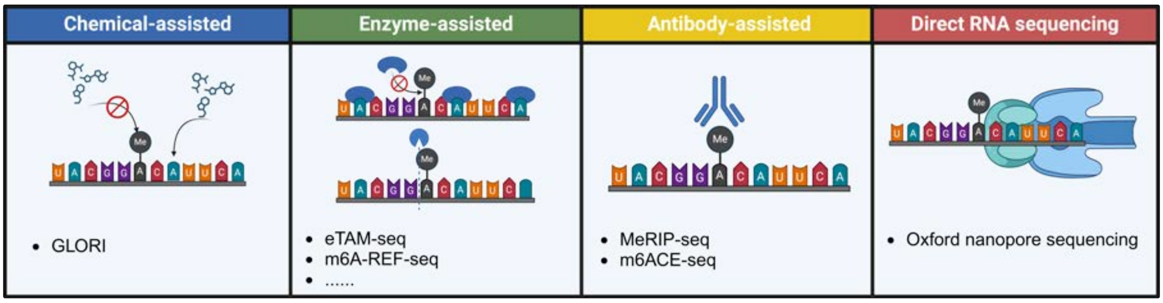


Four sequencing strategies used by m6A quantitative profiling techniques

- **Chemical-assisted**: Uses specific  chemicals to deaminate normal A sites but keep m6A modification sites intact
- **Enzyme-assisted:** Uses enzymes to capture or cleave at m6A modification sites
- **Antibody-assisted**: Uses the antibody-immunoprecipitation approach to capture RNA fragments with m6A modifications
- **Direct RNA sequencing**: Deciphers the m6A modification information by detecting distinct current shift patterns compared to their unmodified counterparts.

Technqiues in different strategy groups are considered orthogonal technique pairs. One exception can be the techniques in antibody-assisted and direct RNA sequencing strategy groups. They are not considered orthogonal because the computational detection method in direct RNA sequencing techniques incorporates m6ACE-seq sequencing results as its training labels. m6ACE-seq is classified in the antibody-assisted strategy group.

In each orthogonal technique pair, overall and condition-specific sample integration methods can be employed. The overall integration method is the sum of m6A counts and total counts at each site from all samples within each technique. As for the condition-specific integration method, summation takes place on samples with common cell lines or tissues.

The IDR analysis-based integration results are presented in Genomic Ranges and CSV format in the m6AConquer database. For each orthogonal technique pair, the results contain the genomic locations and IDR values of all reproducible sites, in addition to their m6A methylation ratios, posterior probabilities, and Benjamini-Hochberg (BH) corrected p-values as modelled by each technique. Furthermore, a merged Genomic Range and CSV files are provided, containing the union of reproducible sites identified in all orthogonal technique pairs, along with the names and numbers of supported orthogonal technique pairs. Users can access and download these results from the Download page.

### V. Codes

The codes for processing sequencing data is available on GitHub.

The fitting of beta-binomial mixture model and other candidate models can be implemented through `OmixM6A` R package, which is also available on GitHub.
